# A Screening Approach Unveils an Unknown Mn^2+^-dependent Endopolyphosphatase Activity in Yeast

**DOI:** 10.1101/2025.01.19.633793

**Authors:** Sandra Moser, Gloria Hans, Adolfo Saiardi, Samuel Bru, Asli A. Taskin, Chris Meisinger, Henning J. Jessen

## Abstract

Inorganic polyphosphate (polyP) is a ubiquitous biopolymer composed of multiple orthophosphates connected by energy-rich phosphoanhydride bonds. In organisms, polyP is digested by two types of enzymes: exopolyphosphatases, which shorten the chain from the ends by cleaving off monophosphate units, and endopolyphosphatases, which cut the chain internally. While several continuous methods are available to monitor exopolyphosphatase activity, endopolyphosphatase activity assays are less common and typically involve multiple tedious steps. Here, we introduce FRET-polyP_8_, a novel probe for real-time detection of endopolyphosphatase activity. The FRET assay enabled rapid, highly sensitive, single-step detection of specific endopolyphosphatase activity both from isolated proteins and cell extracts. The simple read-out additionally enabled enzyme inhibitor screening. Furthermore, a novel Mn^2+^-dependent endopolyphosphatase activity in baker’s yeast was detected in a quadruple mutant, highlighting the ability to screen for metal-dependence of new endopolyphosphatase activity. This approach thus represents a significant addition to existing methodologies, facilitating the discovery and classification of new endopolyphosphatases and their inhibitors to advance our understanding of polyP metabolism and regulation.

## Introduction

Inorganic polyphosphate (polyP), a linear polymer composed of multiple condensed orthophosphates, found in all organisms, is present in a large variety of different chain lengths (from 3 to several 1000).^[1]^ Despite its ubiquity, polyP research has received far less attention than more extensively studied biopolymers such as DNA/RNA and peptides. As a result, there is still much to be explored in terms of biological pathways, function, regulation, enzymology, chemical synthesis, and probe development.^[2]^ In recent years, however, the synthesis of modified polyphosphates of known chain-length has been developed^[3]^ and significant progress has been made in revealing the metabolism in prokaryotes and yeast^[4]^, as well as developing an understanding of the diverse functions of polyP.^[5]^ Research into polyP functions also requires knowledge about and study of enzymes involved in its metabolism.^[4,6]^

The first enzyme responsible for the synthesis of polyP in *E. coli*, the polyP kinase PPK, was identified by Arthur Kornberg in 1956.^[7]^ Nowadays referred to as PPK1, this enzyme catalyzes the transfer of the terminal phosphate group from ATP to polyP.^[8]^ One year later, Kornberg found that this reaction is reversible, degrading polyP to generate ATP from ADP. Another enzyme that degrades polyP by cleaving the terminal phosphate group but releasing free monophosphate (P_i_), is the exopolyphosphatase PPX, also discovered by Kornberg in 1993.^[9]^ In the following year he isolated the homolog of PPX in *Saccharomyces cerevisiae*, known as scPPX1.^[10]^ Unlike bacteria, organisms ranging from archaea to mammals also host a strong endopolyphosphatase (PPN) activity, which cleaves internal phosphoanhydride bonds.^[11]^ In yeast, three endopolyphosphatases have been identified: 1) the vacuolar PPN1, which exhibits both exo- and endopolyphosphatase activity, 2) the vacuolar PPN2 and 3) the cytosolic diadenosine and diphosphoinositol polyphosphate phosphohydrolase (DDP1).^[12]^ The four human homologs of DDP1, belonging to the diphosphoinositol polyphosphate phosphohydrolases (DIPPs) subfamily of NUDIX (nucleoside diphosphate-linked moiety X) hydrolases – DIPP1 (NUDT3), DIPP2 (NUDT4) and DIPP3a/b (NUDT10 and NUDT11) – also exhibit endopolyphosphatase activity. Among these, NUDT3 has the highest activity to degrade polyP in the presence of Zn^2+^.^[13]^ A state-of-the-art method for determining polyphosphatase activity is polyacrylamide gel electrophoresis (PAGE).^[14]^ Exopolyphosphatase activity continuously shortens the polyP chain. Consequently, complete degradation of polyP by exopolyphosphatase results in the disappearance of gel bands, which correlates with an increase in released monophosphate (P_i_) quantified using the colorimetric malachite green assay.^[15]^ In contrast, endopolyphosphatase activity increases the toluidine blue or DAPI staining intensity of small polyP fragments, which remain visible on the gel. ^[12d,13a,16]^ Additionally, PAGE can confirm polyphosphatase inhibition, as the intact long-chain polyP substrate remains detectable when enzymatic activity is fully suppressed by the addition of inhibitors.^[12d,13a]^ However, these PAGE approaches are labor-intensive and only the colorimetric P_i_ detection of exopolyphosphatase activity offers continuous monitoring.^[17]^ The only high-throughput method for detecting both exo- and endopolyphosphatase activity with polyP as substrate uses chromophore- or fluorophore-labeled polyP, allowing reaction tracking via UV/Vis or fluorescence. Nevertheless, endopolyphosphatase activity determination requires a two-step process: first converting bis-labeled polyP with endopolyphosphatase into single-labeled fragments, then monitoring chain shortening by an exopolyphosphatase.^[18]^ Continuous high-throughput inhibitor screening methods have been reported for assays with nucleoside diphosphate-linked moiety X-substrates, as for example diadenosine polyphosphates (Ap_n_A), which can be understood as nucleoside-terminated short-chain polyP. These methods involve a fluorescently labeled *E. coli* phosphate binding protein (PBP) as P_i_-sensor^[19]^, coupling ATP release to luciferase luminescence or the hexokinase reaction^[20]^, or using a substrate capable of Förster resonance energy transfer (FRET).^[21]^

Here, we present the chemical synthesis and application of the first polyP_8_ derivative labeled with a FRET pair at the chain termini. This new probe enables real-time and single-step detection of endopolyphosphatase activity as well as enzyme inhibition screening. Designed for use with a plate reader, the assay is fast, highly sensitive and simple, making it in principle compatible with high-throughput screening approaches. We demonstrate the probe’s specificity for detecting endopolyphosphatase activity with isolated proteins and confirm its utility in identifying and validating DDP1 inhibitors. Additionally, we show that this probe can monitor endopolyphosphatase activity continuously in cell extracts, enabling us to discover a novel Mn^2+^-dependent strong endopolyphosphatase activity in a yeast quadruple mutant that was believed to be devoid of any remaining polyP degrading activity.^[22]^

## Results and Discussion

### Design and Synthesis of the FRET-Probe

Commercially available polyphosphates are mixtures with variable chain lengths, complicating analytical techniques like NMR and mass spectrometry and more importantly, limit modifications to symmetric end-groups or mixtures of un-, mono- and bis-labeled products.^[23]^ In contrast, the chemical bottom-up synthesis of a defined short-chain polyP^[3,24]^ offers significant advantages by enabling precise analytical characterization and allowing the introduction of different modifications at the termini with full control. Attaching a FRET donor and FRET acceptor dye to the ends of polyP allows monitoring of its structural integrity. When the polyP is intact, internal Förster resonance energy transfer occurs. Cleavage of an internal phosphoanhydride bond will disrupt the energy transfer. Consequently, a FRET labeled polyP would be an efficient tool to detect endopolyphosphatase activity by monitoring FRET ratios (Scheme 1a). Cy3/Cy5 is a well-established donor/acceptor FRET pair.^[25]^ Their sulfonate analogs are water-soluble^[26]^ and commercially available with different moieties for attachment, such as azides or NHS esters. The synthesis of the sulfo-Cy3 and sulfo-Cy5 modified polyP with a defined chain length of eight phosphate units is based on an iterative polyphosphorylation using the cyclic pyrophosphoryl P-amidite reagent (**1**) (c-PyPA) (Scheme 1b).^[27]^ In the first round, c-PyPA (**1**) was activated with dicyanoimidazole (DCI) and coupled on one side selectively to the tetrabutylammonium (TBA) salt of pyrophosphate (**2**), a form in which PP_i_ is soluble in organic solvent. After oxidizing P^III^ to P^V^, the cyclotriphosphate intermediate^[28]^ was linearized with propargylamine, thereby introducing a terminal alkyne as clickable handle. This entire sequence was carried out as a one-pot reaction. The resulting propargylamido-polyP_5_ (**3**) was converted into its TBA salt, enabling a second tripolyphosphorylation step. Here, diaminopropane was used as nucleophile allowing orthogonal functionalization of the obtained polyP_8_ **4** on the termini. Diaminopropane prevents chain shortening via cyclization and anhydride cleavage, unlike ethylenediamine, which can form a phosphorus diamidate thus removing a terminal phosphate unit.^[29]^ The FRET dyes were then attached by amide bond formation and azide-alkyne cycloaddition. Amide coupling was challenging due to rapid hydrolysis of the sulfo-Cy3-NHS ester under the conditions applied. The overall yield of the click reaction was reduced by compound loss during multiple washing and precipitation steps, explaining the comparably low yields.

Residual copper ions were removed using Chelex® resin. All isolated compounds were purified by strong anion exchange (SAX) chromatography with NH_4_HCO_3_ or NaClO_4_ as elution buffer, resulting in the ammonium or sodium salts. A defined chain-length of eight phosphates of the FRET-polyP_8_ (**6**) was confirmed by ^31^P{^1^H}-NMR spectroscopy (Scheme 1c). The two terminal phosphoramidates have characteristic shifts between 0 and

-2 ppm and the internal phosphoranhydrides resonate at around −22 ppm with a ratio of 1:1:6 (for a ^31^P-NMR chemical shift table for condensed phosphates, see Accounts Chem. Res.^[24a]^). Assignment of the phosphoramidates was performed during analysis of compound **4** using the ^1^H-^31^P-HMBC-NMR pulse sequence (Figure SI-1), which is also applicable to compound **6** (Figure SI-2). High purity of **5** and **6** was verified by UPLC-MS analysis (Figures SI-3 and SI-4). The bottom-up synthesis enabled isolation of fully and asymmetrically end-labeled polyP with known chain-length, eliminating the need for a digestion-reaction-dependent determination of end-labeling efficiency. This would be typically required when using 1-ethyl-3-(3-(dimethylamino)-propyl)carbodiimide (EDAC) mediated coupling reactions to obtain symmetrically modified polyPs^[18]^. Furthermore, the use of a single compound rather than a substrate mixture simplified analysis by techniques such as NMR, UPLC and mass spectrometry. Fluorescence spectroscopy demonstrated an efficient internal energy transfer within the FRET polyP_8_ (**6**) (Scheme 1d,e). Excitation of sulfo-Cy3 at 500 nm resulted mainly in the fluorescence emission spectrum of sulfo-Cy5.

## Determination of Endopolyphosphatase Activity with Isolated Proteins

Previously, PAGE studies showed that blocking of both termini in polyphosphates protects them from degradation by exopolyphosphatases, while they remain a substrate for endopolyphosphatases.^[3,18]^ Consistent with these findings, the FRET-polyP_8_ resisted degradation by the exopolyphosphatase PPX from *E. coli* as evidenced by its sustained high FRET ratio, which represents the ratio between the emission intensity of acceptor dye Cy5 at 670 nm and the total emission intensity of Cy5 at 670 nm and the donor dye Cy3 at 565 nm (Eq. 1). In contrast, treatment with the endopolyphosphatase DDP1 from *Saccharomyces cerevisiae* efficiently cleaved FRET-polyP_8_ (**6**), time-dependently disrupting the energy transfer and thus resulting primarily in the Cy3 emission spectrum with a low FRET ratio (Figure 1a,c). Hence, the FRET-polyP_8_ probe serves as a tool for time-resolved detection of endopolyphosphatase activity.

**Figure 1.**
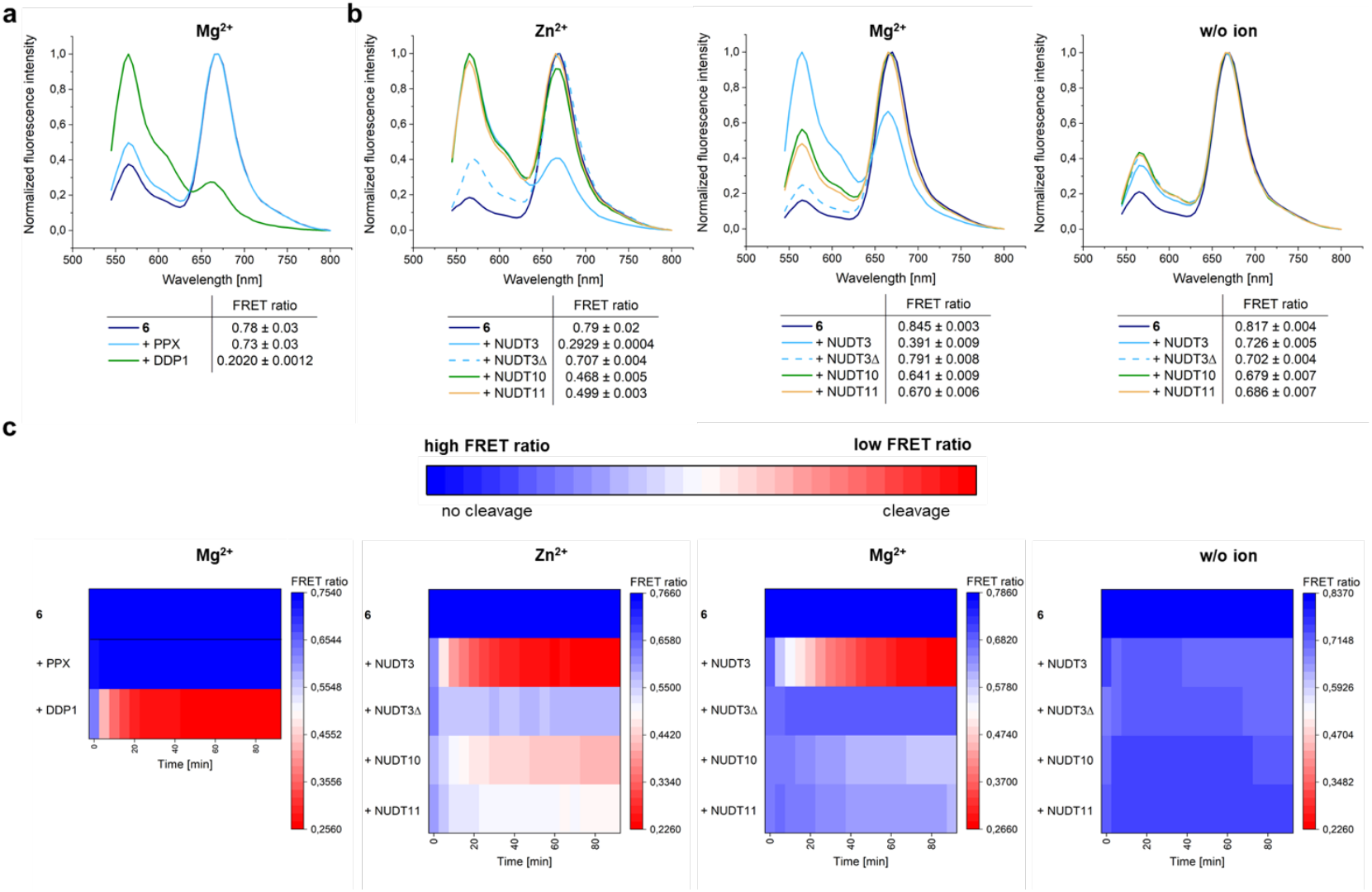
Endopolyphosphatase assay with isolated enzymes. **a**, Normalized fluorescence intensity and FRET ratios 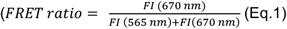, FI: fluorescence intensity) of **6** (100 nM in 1× reaction buffer) after 30 min incubation at 37 °C with either the exopolyphosphatase PPX (2 μg) or the endopolyphosphatase DDP1 (2 μg) or without enzyme (control) in presence of 10 mM Mg^2+^. Excitation at λ = 500 nm. FRET-ratios are shown as mean ± SD (*n* = 3, SD: standard derivation). **6** was only cleaved by the endopolyphosphatase. **b**, Normalized fluorescence intensity spectra and FRET ratios of **6** (100 nM in 1× reaction buffer) after 30 min incubation at 37 °C with NUDIX enzymes (2 μg) and addition of 5 mM ZnSO4 or MgSO4 or without additional divalent ion. Excitation at λ = 500 nm. NUDT3 was showing the highest endopolyphosphatase activity in presence of Zn^2+^, in accordance with literature results.^[13b]^ **c**, Time courses of the FRET ratio change over 90 min with the same reaction conditions as described in **a** and **b.** DDP1 and NUDT3 in presence of divalent cations were cleaving the fastest. Buffer compositions are listed in Table SI-3.

To further validate this, three of the human homologs of DDP1 were also tested (Figure 1b), for which metal-dependent substrate specificity has been shown. In accordance with earlier studies of Samper-Martín *et al*.^[13b]^, NUDT3 demonstrated high endopoly-phosphatase activity, with much greater activity in the presence of Zn^2+^ compared to Mg^2+^. As expected, neither the inactive NUDT3 mutant with a replacement of the catalytic glutamic acid at position 70 to alanine (NUDT3Δ = NUDT3E70A) nor the absence of the metal cofactor resulted in any detectable activity. NUDT10 and NUDT11, identified as endopolyphosphatases with lower activity than NUDT3^[13b]^, also exhibited lower activity in the presence of 5 mM Zn^2+^ compared to NUDT3 with either Zn^2+^ or Mg^2+^. Additionally, it was observed that NUDT10 and NUDT11 showed minimal activity when Mg^2+^ was used as cofactor and that there was no activity without any ion. The time course of the FRET ratio change upon treatment of FRET-polyP_8_ (**6**) with the different endopolyphosphatases is displayed in Figure 1c as a heat-map. Among the tested enzymes, DDP1 with Mg^2+^ as cofactor exhibited the highest activity, achieving complete cleavage within 20 min. NUDT3 fully decomposed FRET-polyP_8_ (**6**) in 30 min in the presence of Zn^2+^ and in 50 min with Mg^2+^. NUDT10 and NUDT11 did not fully digest the probe within 90 min under any tested conditions, however partial cleavage was observed. All these results support the conclusion that this assay accurately recapitulates the described activity of the NUDIX isoforms and therefore our FRET-assay offers a simple and highly sensitive method for analyzing endopolyphosphatase activity. Unlike PAGE-based techniques, less procedural steps and lower concentrations of polyP are needed. While electrophoresis analysis typically requires 200-250 nmol polyP^[16,13a]^, fluorescence spectroscopy works with 0.75 pmol of the FRET-polyP_8_, all in P_i_ units, corresponding to nanomolar probe concentrations. Moreover, PAGE analyses are typically run overnight and require staining and destaining procedures. Similar to the method developed by Hebbard *et al*.^[18]^, this assay can in principle be adapted to high-throughput screenings of active enzymes.

### Inhibitor Screening of DDP1

Since this FRET assay is easy to implement using a plate reader and simple to analyze via FRET ratio calculations, it can be further used to screen endopolyphosphatase inhibitors (Figure 2a). For validation, we tested known DDP1 inhibitors.^[13a]^ However, the exact IC_50_ values for these inhibitors have not previously been reported, only their ability to inhibit cleavage. In addition, we screened other divalent cations as potential inhibitors. To ensure that the known and potential inhibitors themselves do not degrade the FRET probe, we first investigated the probe’s stability against increasing concentrations of each potential inhibitor by monitoring changes in fluorescence spectra (Figure SI-5) or variations in the FRET ratio (Figure 2b). FRET-polyP_8_ (**6**) was very stable in presence of MgCl_2_, Heparin and CaCl_2_. Slight decomposition was observed at concentrations starting from 25 mM of NaF and ZnSO_4_ and at 100 mM of MnSO_4_. However, only 0.3 mM FeSO_4_ was sufficient to completely reduce the FRET ratio, by causing precipitation of **6**, leading us to exclude FeSO_4_ from further inhibitor screening (Figure 2c and Figure SI-6). Full inhibition of DDP1 activity was achieved with 1 μM heparin and 100 mM CaCl_2_, NaF and ZnSO_4_. It is important to note that the addition of 100 mM NaF and ZnSO_4_ alone reduced the FRET ratio of the FRET-polyP8 from ∼ 0.7 to ∼ 0.5 (see Figure 2b), defining ∼ 0.5 as the maximum observable inhibition level and indicating complete enzyme inhibition at this ratio. MgCl_2_, which did not degrade the FRET probe, achieved only partial inhibition at high concentrations while MnSO_4_ showed no inhibition within the tested concentration range. These findings align well with literature,^[13a]^ except for MnSO_4_’s lack of inhibition. Lonetti *et al*.^[13a]^ observed DDP1 inhibition with 2 mM MnCl_2_, but at a 200-fold lower amount of DDP1, potentially accounting for the different results. Among the tested inhibitors, only CaCl_2_ and heparin followed a dose-response-curve, enabling IC_50_ determination (Figure 2d). Heparin, with a delineated IC_50_ value in the nanomolar range, proved to be significantly more potent than CaCl_2_, which required millimolar concentrations for effective inhibition. This is the first time that such values have become readily available in polyP research.

**Figure 2.**
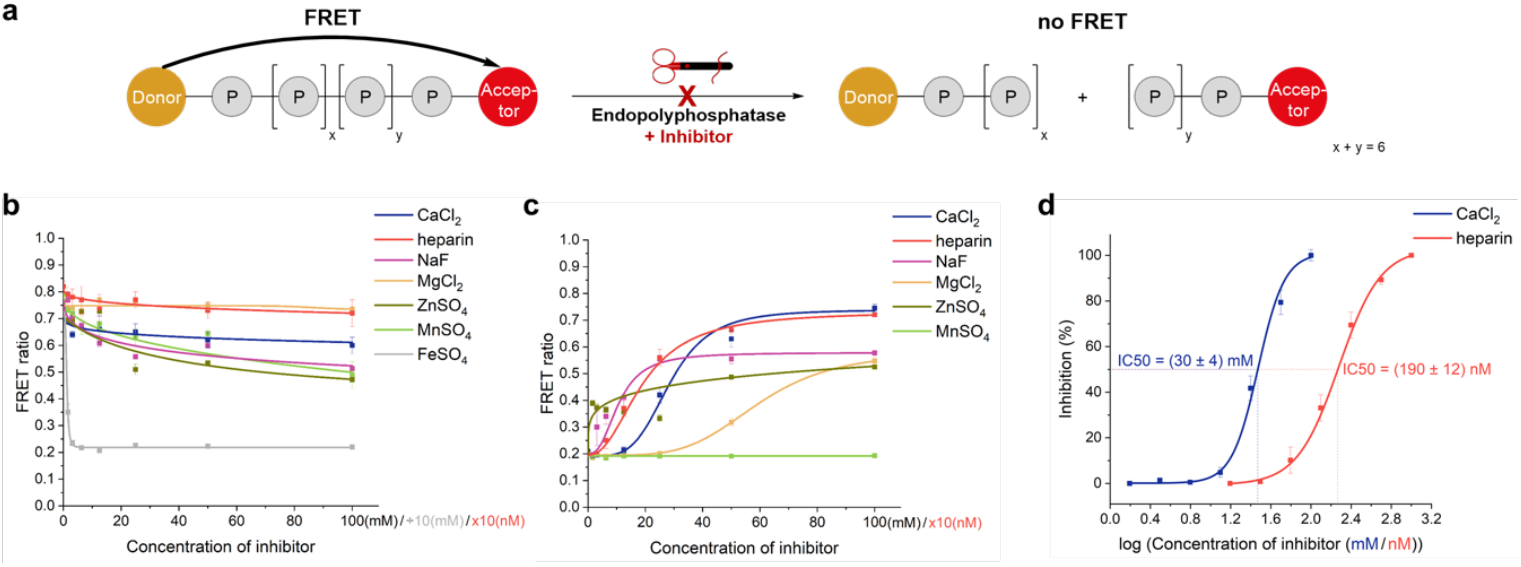
DDP1 inhibition screening. **a**, Graphical representation of the inhibition assay principle: endopolyphosphatase inhibition prevents polyP cleavage, sustaining the FRET signal. **b**, Stability control of **6** in the presence of different potential DDP1 inhibitors: FRET ratio changes of **6** (100 nM) after incubation with different salt concentrations at 37 °C after 30 min (heparin sodium salt: 0-1,000 nM (calculated with average molecular weight of 17 kDa), FeSO4: 0-10 mM, others: 0-100 mM, all in addition to 10 mM MgSO4 as part of the reaction buffer, listed in Table SI-3). FRET ratios are shown as mean ± SD (*n* = 3). **c**, DDP1 inhibition screening: FRET ratio changes of **6** (100 nM in 1× reaction buffer) after incubation with DDP1 (2 μg) at 37 °C for 30 min in the presence of different potential inhibitors, tested across different concentrations (heparin in nM, all others in mM). **d**, Dose-response curves and IC50 analysis of inhibitors of DDP1. Normalized inhibitions were calculated using 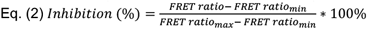. Error bars were calculated with Gauss’ law of error propagation. A non-linear curve fitting for dose response curves was performed to determine IC50 values and their standard deviation.

### Discovery of a New Endopolyphosphatase Activity in Yeast Extracts

Determination of endopolyphosphatase activity with FRET-polyP_8_ (**6**) was also tested in *Saccharomyces cerevisiae* cell extracts. We examined wild-type (WT) as well as knockout *S. cerevisiae* strains lacking some or all known yeast endopolyphosphatases and other enzymes involved in yeast phosphate homeostasis (Figure 3a).^[22,30]^ In the presence of 10 mM Mg^2+^, the WT strain showed very fast cleavage of **6** (within ca. 10 minutes), as expected. In the quadruple KO strain *ppx1Δ ppn1Δ ppn2Δ vtc4Δ*, missing two of the three known yeast endopolyphosphatases and lacking endogenous polyP (since VTC4 is the polyP synthase in yeast)^[31]^, a slower decline in the FRET ratio was observed. We attribute this reduced cleavage rate to DDP1, being the only remaining endopolyphosphatase activated with Mg^2+^. In contrast, the triple knockout strain *ppx1Δ ppn1Δ ppn2Δ*, which contains high levels of endogenous long-chain polyP, did not show efficient cleavage of the probe. We hypothesize that DDP1 will be predominantly engaged in cleaving the very abundant endogenous polyP rather than the FRET probe. As anticipated, no distinct endopolyphosphatase activity was observed in the strain *ppx1Δ ppn1Δ ppn2Δ ddp1Δ*, in which all known yeast endopolyphosphatases are knocked out. However, since this strain also contains endogenous polyP, it is again possible that any other weakly active endopolyphosphatase, if present, might preferentially target endogenous polyP, thereby escaping detection. Interestingly, the addition of 10 mM MnSO_4_, resulted in a clear reduction of the FRET ratio in all four yeast extracts (Figure 3b). The decrease in cleavage velocity correlates with the number of endopolyphosphatase knockouts. Also, cleavage is much faster in the absence of endogenous polyP in VTC KO strains. A control experiment with FRET-polyP_8_ (**6**) in lysis buffer, supplemented with 10 mM MnSO_4_, demonstrated that the decrease in FRET ratio was not due to decomposition of FRET probe **6** by MnSO_4_ or by precipitation. To further confirm that the observed effect was not related to FRET-polyP_8_ (**6**) itself, we investigated the impact of Mg^2+^ and Mn^2+^ on cellular polyP. PAGE analysis (Figure 3c) of the digested polyP product with WT *S. cerevisiae* extracts, either without the addition of a divalent cation or with 10 mM Mg^2+^, showed no remaining polyP, as the endogenous polyP was cleaved by both endo- and exopolyphosphatases. When Mn^2+^ was added, some bands migrating as small-chain polyP became detectable. This may be due to Mn^2+^ slowing down the PPX digestion. In contrast, the quadruple mutant *ppx1Δ ppn1Δ ppn2Δ ddp1Δ* samples retained bands on PAGE corresponding to high molecular weight polyP in the absence of salt or with Mg^2+^, but showed efficient cleavage of the high polymeric polyP into smaller polyP chains in the presence of Mn^2+^, clearly demonstrating endopolyphosphatase activity.

**Figure 3.**
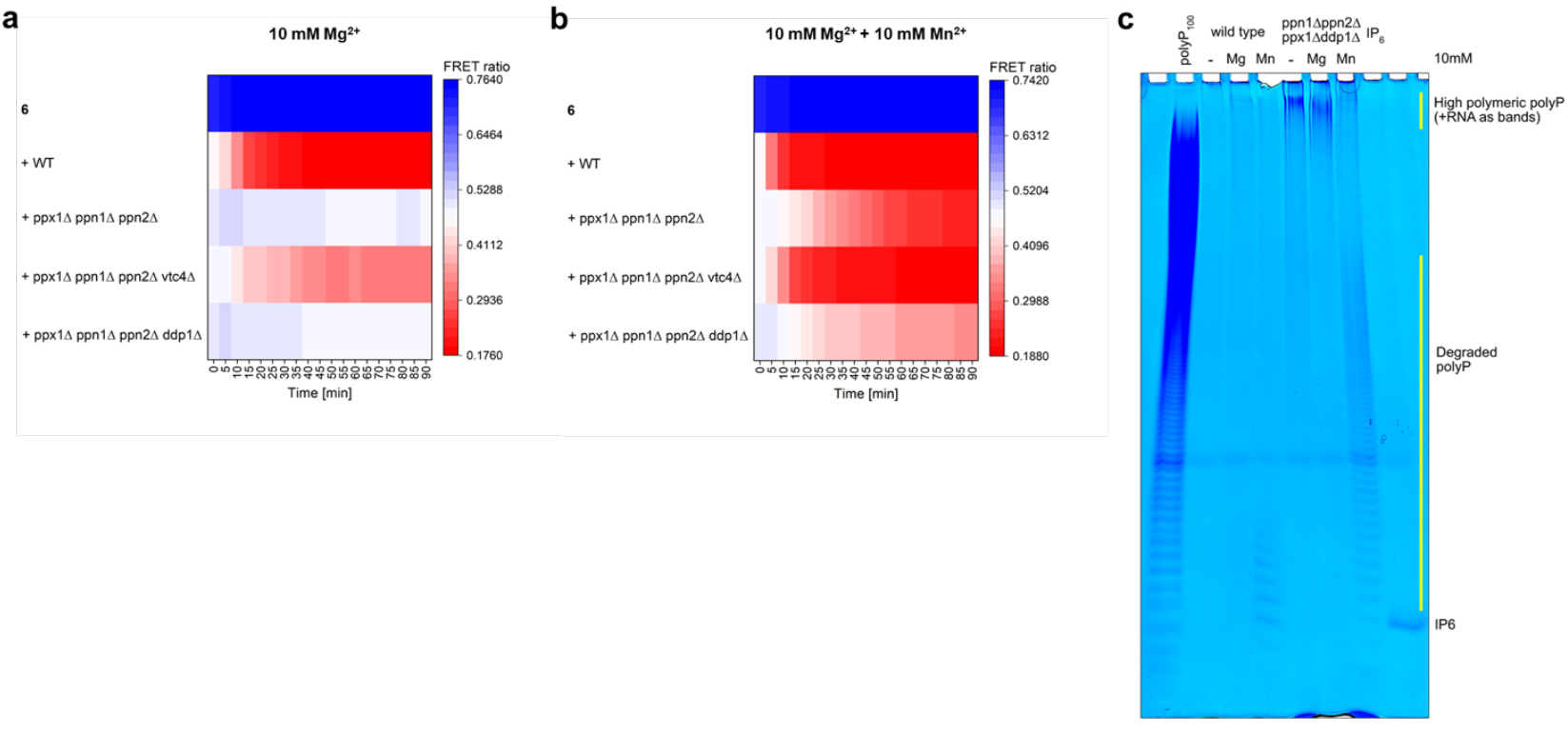
Influence of manganese ions versus magnesium ions on endopolyphosphatase activity in baker’s yeast extracts. **a**, Monitoring of the FRET ratio changes over 90 min during incubation of **6** (100 nM) with yeast extracts from different mutant strains (1 mg) in the presence of 10 mM MgSO4. Knock-out mutants are as follows: *ppx1Δ* (exopolyphosphatase), *ppn1Δ* and *ppn2Δ* (endopolyphosphatases), *ddp1Δ* (diadenosine and diphosphoinositol polyphosphate phosphohydrolase). **b**, FRET ratio changes in the presence of additional 10 mM MnSO4, showing endopolyphosphatase activity also in the KO strains that had no significant cleavage of **6** in the absence of MnSO4. **c**, Influence of Mg^2+^ (10 mM) and Mn^2+^ (10 mM) on cellular polyP in WT yeast and the quadruple mutant. Extracts were incubated at 37 °C for 60 min and polyP was visualized on a 30% polyacrylamide gel.

Taken together, the findings from the PAGE analysis of cellular polyP in the presence of Mn^2+^, enabled by the FRET-polyP_8_ assay, suggest the existence of another, so far unknown, endopolyphosphatase in *S. cerevisiae*. This activity is hidden under “normal” conditions but becomes apparent when activated with Mn^2+^. The distinct cleavage observed in the FRET assay, even in strains containing endogenous polyP, is pointing to high enzymatic activity under these new conditions. Manganese plays a crucial role as cofactor in a wide variety of metalloproteins. It is important to note that yeast cells normally accumulate manganese at concentrations ranging from 0.04 to 2.0 mM in the cytosol.^[32]^ Therefore the addition of 10 mM MnSO_4_ appears to be above the physiological range. However, such ions can be transported, stored and accumulated in the yeast vacuole to higher concentrations of 14 mM^[33]^; of note, yeast endopolyphosphatases PPN1, PPN2 are both highly enriched in the vacuole, which is also the main polyP storage organelle.^[34]^ Thus, the identified activity may well play an overlooked role in vacuolar polyP homeostasis. The identification of the enzyme(s) responsible for this activity remains as the next task. In this context, the FRET-polyP_8_ (**6**) can be used to develop a biochemical purification protocol to isolate the activity.

## Conclusion

In conclusion, we developed the first one-step continuous endopolyphosphatase activity assay, using a FRET-labeled polyP substrate **6** with a defined chain-length of eight phosphates. This is the first example of an asymmetrically modified FRET-polyP; orthogonal terminal modifications (alkyne, amine) now offer the possibility of developing new polyP tools, such as combined photoaffinity or nonhydrolyzable pulldown probes with differently modified end-groups. This is important given that many enzymes involved in mammalian polyP metabolism are likely still unidentified.^[35]^ The FRET-labeled polyP_8_ **6** can be used in a FRET assay in a plate reader to sensitively and in real-time determine endopolyphosphatase activity and endopoly-phosphatase inhibition with a simple read-out. By applying the FRET assay to known endopolyphosphatases and inhibitors, we successfully validated its functionality. This assay is well-suited for high-throughput screenings of active enzymes and a high-throughput inhibitor screening, enabling fast detection of enzymatic activity or inhibition. The assay proved to be particularly convenient for testing the effect of the presence of different metal ions on enzyme activity. Additionally, we demonstrated its compatibility with yeast cell extracts. We observed significant digestion of **6** with yeast extracts derived from a quadruple mutant lacking all known yeast polyphosphatases in the presence of Mn^2+^. Manganese restores endopolyphosphatase activity within the yeast cell extract which was also shown by PAGE on cellular polyP. FRET-polyP_8_ (**6**) thus enabled the discovery of a new manganese-ion-activated yeast endopoly-phosphatase activity that appears to be highly efficient. This tool also holds promise to isolate the unidentified Mn^2+^-dependent endopolyphosphatase in yeast, whose existence can now be postulated based on the findings reported herein. Furthermore, this novel tool can also be applied to other areas of polyP research, such as studying its subcellular localization after transporter-mediated uptake. Unlike symmetrically fluorescently labeled polyP uptake studies^[23c]^, the use of FRET-polyP_8_ (**6**) will offer distinct advantages of monitoring its integrity during uptake and distribution, allowing to distinguish whether the fluorescent signal inside the cell results from the intact compound or from the fluorophore itself due to decomposition. In addition, FRET-polyP_8_ (**6**) can be used to investigate, whether lysine polyphos-phorylation is a covalent post translational modification (no FRET signal) or an ionic interaction (FRET signal), contributing to the ongoing debate.^[36]^

## Supporting Information

Supporting information include supporting figures, experimental procedures and characterization data. The authors have cited additional references within the Supporting Information.^[37]^

## Acknowledgements

We appreciate the support of G. Liu, M. Lu, I. Prucker and A. Shukla from the Jessen group for CE and HRMS measurements. We also thank Dr. S. Braukmüller and C. Warth from the Analytical Service Team of the University of Freiburg for NMR and HRMS measurements, respectively. Additionally, we would like to thank Prof. Dr. Josep Clotet for valuable discussions. This work has received funding by the Deutsche Forschungsgemeinschaft (DFG, German Research Foundation, project number 445698446) in collaboration with the Indian Department of Biotechnology (DBT). This research was supported by the Deutsche Forschungsgemeinschaft (DFG) under Germany’s Excellence Strategy (CIBSS-EXC-2189-Project ID 390939984, to HJJ). A. S. was supported by the UKRI Medical Research Council grant MR/T028904/1.

---

**Scheme 1.**
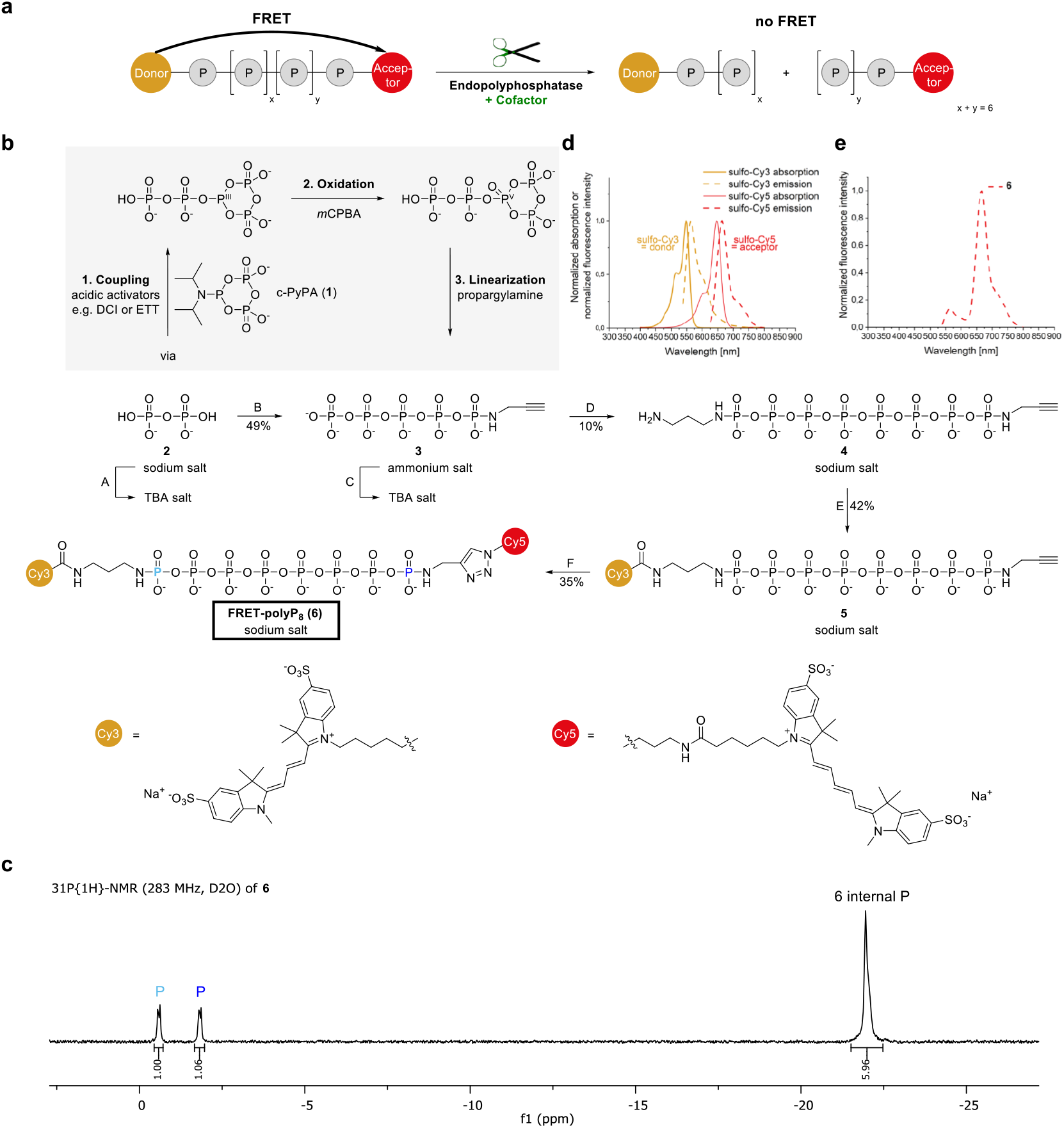
Design, synthesis and characterization of **6. a**, Design of a FRET probe for detecting endopolyphosphatase activity. **b**, Synthesis of **6**: A, Dowex® 50WX8 H^+^ column followed by neutralization with TBAOH. B, 1. DCI (3.0 eq.), c-PyPA (**1**) (0.075 M in MeCN, 1.3 eq.), −20 °C, 15 min. 2. *m*CPBA (1.5 eq.), −20 °C 0 °C, 20 min. 3. Propargylamine (5.0 eq.), 0 °C r.t., 60 min. C, Chelex® 100 column preconditioned with TBA(Br) (500 mM). D, 1. ETT (11 eq.), **1** (0.075 M in MeCN, 3.2 eq.), DMF, r.t., 45 min. 2. *m*CPBA (4.7 eq.), −15 °C, 40 min. 3. 1,3-Diaminopropane (40 eq.), 0 °C r.t., 30 min. E, Sulfo-cyanin3-NHS-ester (2.0 eq.), 0.1 M NaHCO3, r.t., 3.5 h. F, Sulfo-cyanin-5-azide (1.5 eq.), CuSO_4_ × 5 H_2_O (20 mM in H_2_O, 0.5 eq.), THPTA (50 mM in H_2_O, 2.5 eq.), sodium ascorbate (10.0 eq.), r.t., 3.5 h. **c**, ^31^P{^1^H}-NMR spectrum of **6** confirming a defined chain-length of eight phosphates. **d**, Normalized absorption (10 μM) and normalized fluorescence intensity (100 nM) spectra of sulfo-Cy3 (excitation at λ = 500 nm) and sulfo-Cy5 (excitation at λ = 590 nm) in water. **e**, Normalized fluorescence intensity spectrum of **6** in water (100 nM in Cy3 terms, not Pi units (more details in Table SI-1), excitation at λ = 500 nm) demonstrating effective FRET. Abbreviations: TBA: tetrabutylammonium, DCI: 4,5-dicyanoimidazole, *m*CPBA: *meta*-chloroperbenzoic acid, ETT: 5-(ethylthio)-1*H*-tetrazole, THPTA: tris[(1-hydroxy-propyl-1*H*-1,2,3-triazol-4-yl)methyl]amin.

