## Supplement for "A Screening Approach Unveils an Unknown Mn^2+^-dependent Endopolyphosphatase Activity in Yeast"

**Abbreviations:**

| | |
|---|---|
| **BSA** | bovine serium albumin |
| **CE** | capillary electrophoresis |
| **Cy3** | cyanine3 |
| **Cy5** | cyanine5 |
| **DCI** | 4,5-dicyanoimidazole |
| **DMF** | dimethylformamide |
| **DTT** | dithiothreitol |
| **ESI** | electrospray ionization |
| **ETT** | 5-(ethylthio)-1*H*-tetrazole |
| **HEPES** | 4-(2-hydroxyethyl)piperazine-1-ethanesulfonic acid |
| **HRMS** | high resolution mass spectrometry |
| ***m*CPBA** | *meta*-chloroperbenzoic acid |
| **MS** | mass spectrometry |
| **m/z** | mass-to-charge ratio |
| **NHS** | N-hydroxysuccinimide |
| **NMR** | nuclear magnetic resonance |
| **PMSF** | phenylmethylsulfonyl fluoride |
| **$PP_i$** | pyrophosphate |
| **qTOF** | quadrupole time-of-flight |
| ***S. cerevisiae*** | *Saccharomyces cerevisiae* |
| **TBA** | tetrabutylammonium |
| **TFA** | trifluoroacetic acid |
| **THPTA** | tris[(1-hydroxy-propyl-1*H*-1,2,3-triazol-4-yl)methyl]amine |
| **TRIS** | tris(hydroxymethyl)aminomethane |
| **UPLC** | ultra-performance liquid chromatography |

### 1. Synthesis of FRET-polyP$_8$

#### 1.1 General Methods and Materials

Reactions were performed in flame-dried glassware under inert gas atmosphere unless the solvent was water/buffer. Water was purified with a Milli-Q® lab water system. Reagents were purchased from commercial suppliers and were used without further purification. Sulfo-Cy3-NHS-ester and sulfo-Cy5-azide were purchased from Lumiprobe. Solvents were obtained in analytical grade and were used as received. Reaction control was done by $^{31}P\{^1H\}$-NMR.

**Strong anion exchange chromatography** was performed using an automated ÄKTA pure™ system and QSepharose® Fast Flow (Sigma-Aldrich). Crude products were loaded as aqueous solutions and eluted using increasing concentrations of either $NH_4HCO_3$ or $NaClO_4$ solutions.

**Cation exchange** for the preparation of TBA salts was performed with Dowex® 50WX8 $H^+$, followed by neutralization with TBA hydroxide and subsequent lyophilization or with a Chelex® 100 column preconditioned with TBA(Br) (500 mM).

**Lyophilization** was performed using a Christ Alpha 1-4 LDplus.

**Centrifugation** was performed with an Eppendorf 5804R.

**NMR-spectroscopy:** The $^{1}H$-, $^{13}C$-, $^{31}P$-NMR spectra were measured on a Bruker Avance Neo 400 MHz (101 MHz for $^{13}C$, 162 MHz for $^{31}P$) NMR spectrometer with broadband CryoProbe Prodigy and Bruker Avance Neo 700 MHz (176 MHz for $^{13}C$, 283 MHz for $^{31}P$) NMR spectrometer with broadband CryoProbe Prodigy. All signals were referred to an internal solvent signal ($^{1}H$-NMR: $D_2O$: $\delta = 4.79$ ppm). The signals of $^{31}P$-NMR and $^{13}C$-NMR spectra were referenced to an external standard. The chemical shifts are quoted in ppm. The splitting patterns are labeled as: singlet (s), doublet (d), triplet (t), multiplet (m). The coupling constants *J* are given in Hertz (Hz). The evaluation of NMR-spectra was done using the software MestreNova from Mestrelab Research.

**CE-ESI-MS** experiments were performed on a bare-fused silica capillary, activated for 10 min with NaOH (1 M) before first measurement, with a length of 100 cm (50 µm internal diameter and 365 µm outer diameter) on an Agilent 7100 capillary electrophoresis system coupled to a qTOF (6520, Agilent) equipped with a commercial CE-MS adapter and sprayer kit from Agilent. 35 mM ammonium acetate titrated by ammonia solution to pH 9.75 was background electrolyte. Samples were injected by applying 100 mbar pressure for 10 s, followed by an injection of a background electrolyte plug by applying 50 mbar for 5 s. For each analysis, a constant CE current of 23 µA was established by applying 30 kV over the capillary. The sheath liquid was composed of a water-isopropanol (1:1) mixture spiked with mass references. It was introduced at a constant flowrate of 1.5 µL/min. ESI-qTOF-MS was conducted in the negative ionization mode with published settings.[1] Automatic recalibration of each acquired spectrum was performed using reference masses of reference standards (TFA anion, $[M-H]^-$, 112.9855), and (HP-0921, $[M-H+CH_3COOH]^-$, 980.0163. Data were processed using the Agilent CE ChemStation Software.

**UV/Vis absorption for concentration determination** of FRET-poly$P_8$ (**6**) was measured on a Shimadzu UV-1900i UV-Vis spectrometer with quartz SUPRASIL® cuvettes (Ø = 10 mm). The extinction was measured at 548 nm and the concentration was calculated via Beer-Lambert law using ε(Cy3) = 162000 $M^{-1}cm^{-1}$ (Lumiprobe) (Table SI-1).

**Table SI-1:** Measured extinction of FRET-poly$P_8$ (**6**).

| **Compound** | ***λ* / nm[a]** | **A[b]** | **$c_{sample}$ (µM)** | **dilution** | **$c_{stock}$ (µM)** | **solvent** |
|---|---|---|---|---|---|---|
| **FRET-P8** | 548 | 0.539 | 3.327 | 1:200 | 665 | $H_2O$ |

[a] Wavelength of the absorption maximum of sulfo-Cy3. [b] Absorbance.

**Absorption spectra and fluorescence spectra** were recorded on a Tecan Spark® Plate Reader. Time course measurements were run according to the program in Table SI-2.

**Table SI-2:** Plate reader time course settings.

| | | | |
|---|---|---|---|
| **Loop** | Loop type | Duration [hh:mm:ss] | 01:35:00 |
| | Interval type | Fixed [hh:mm:ss] | 00:05:00 |
| | Fluorescence Intensity Scan | Excitation wavelength [nm] | 500 |
| | | Emission wavelength [nm] | 535-800 |
| | | Step size | 5 |
| | Shaking | Duration [s] | 140 |
| | | Mode | Orbital |
| | | Amplitude [mm] | 1 |

#### 1.2 Synthesis of c-PyPA (1)

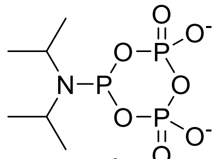

**1**

The compound was synthesized in two steps from pyrophosphate sodium salt as reported previously. Analytical data were identical to literature.[2] The compound was stored as stock solution (0.075 M in MeCN) at -20 °C.

#### 1.3 Synthesis of propargylamido polyP$_5$ (3)

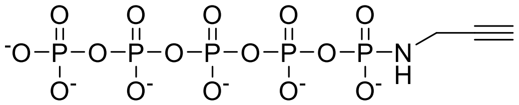

**3**

ammonium salt

According to a reported procedure from J. Singh *et al*.[3]

PP$_i$ × 2 TBA (0.075 M, 20 mL, 1.5 mmol, 1.0 eq.) was added to dry DCI (535 mg, 4.53 mmol, 3.0 eq., dried for one week in desiccator). The mixture was stirred for 5 min before it was cooled down to –20 °C and c-PyPA (0.075 M, 26 mL, 2.0 mmol, 1.3 eq.) was added. The reaction mixture was stirred for 15 min. Then *m*CPBA (70%, 555 mg,

2.25 mmol, 1.5 eq.) was added and the mixture was stirred at –20 °C. After 10 min the cooling bath was removed and the mixture was stirred for further 10 min. Subsequently, propargylamine (480 µL, 413 mg, 7.50 mmol, 5.0 eq.) was added dropwise at 0 °C. After stirring for 60 min at r.t., the crude product was precipitated by cold $NaClO_4$ (0.5 M in acetone, 40 mL), centrifuged, washed with acetone (4×) and purified by strong anion exchange chromatography (Q Sepharose® Fast Flow, increasing concentrations of 1M $NH_4HCO_3$ in $H_2O$). Fractions eluted with 400-500 mM aqueous $NH_4HCO_3$ were combined and lyophilized. To remove buffer, the product was dissolved again in water and lyophilized (2×). The ammonium salt of **3** (411 mg, 738 µmol, 49%) was obtained as a white solid.

Analytical data were identical to literature.[3]

#### 1.4 Synthesis of (3-aminopropyl)amido-propargylamindo-polyP$_8$ (4)

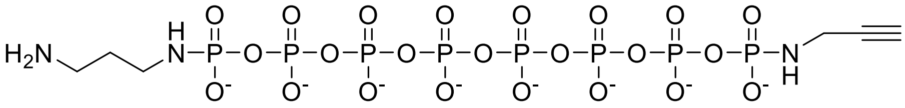

**4**
sodium salt

ETT (18 mg, 0.14 mmol, 11 eq.) and propargylamido-polyP5 × 6 TBA (25 mg, 13 µmol, 1.0 eq.) were both separately coevaporated with dry MeCN (3 × 2 mL). Propargylamido-polyP5 × 6 TBA was dissolved in dry DMF (1 mL) and was added to a solution of ETT in dry DMF (0.5 mL). The mixture was stirred for 10 min before c-PyPA (0.075 M, 0.55 mL, 41 µmol, 3.2 eq.) was added. The reaction mixture was stirred at r.t. for 45 min. Then *m*CPBA (70%, 15 mg, 61 µmol, 4.7 eq.) was added at –15 °C. After 15 min the cooling bath was removed and the mixture was stirred for further 25 min. Subsequently, 1,3-diaminopropane (45 µL, 40 mg, 0.54 mmol, 40 eq.) was added dropwise at 0 °C. After stirring for 10 min at 0 °C the cooling bath was removed and the mixture was stirred for further 20 min. The crude product was precipitated by $NaClO_4$ (0.5 M in acetone, 30 mL), centrifuged, washed with acetone (4×) and purified by strong anion exchange chromatography (Q Sepharose® Fast Flow, increasing concentrations of aqueous $NH_4HCO_3$ in $H_2O$). Product containing fractions were combined and lyophilized. To remove buffer, the product was dissolved again in water and lyophilized (2×) and purified again by strong anion exchange chromatography (High Trap QFF, increasing concentration of aqueous $NaClO_4$-solution (1 M) in $H_2O$). Fractions eluted with 10% aq. $NaClO_4$ were combined, lyophilized, precipitated with cold $NaClO_4$ acetone solution (0.5 M, 35 mL), washed with acetone (3 × 30 mL) and dried to afford **4** (1.2 mg, 1.3 µmol, 10%) as a white solid.

**$^1$H-NMR** (400 MHz, $D_2O$) δ = 3.75 (dd, *J* = 10.6, 2.5 Hz, 2H), 3.17 (t, *J* = 7.1 Hz, 2H), 3.10 (dt, *J* = 10.8, 6.6 Hz, 2H), 2.62 – 2.60 (m, 1H), 1.93 (tt, *J* = 6.9, 6.9 Hz, 2H) ppm. **$^{31}$P{$^1$H}-NMR** (162 MHz, $D_2O$) δ = -1.01 (d, *J* = 20.5 Hz, 1P), -2.28 (d, *J* = 20.0 Hz, 1P), -22.09 – -22.45 (m, 6P) ppm. **$^{31}$P-NMR** (162 MHz, $D_2O$) δ = -1.02 (dt, *J* = 21.0, 10.7 Hz, 1P), -2.28 (dt, *J* = 20.4, 10.5 Hz, 1P), -22.09 – -22.45 (m, 6P) ppm. **$^{13}$C-NMR** (101 MHz, $CDCl_3$) δ = 82.7, 71.5, 38.6, 37.4, 31.1, 28.4 (d, *J* = 7.0 Hz) ppm. **HRMS CE-ESI** calc. for $C_6H_{19}N_3O_{23}P_6^{2-}$ $[M-2H]^{2-}$: 374.4161, found: 374.4173.

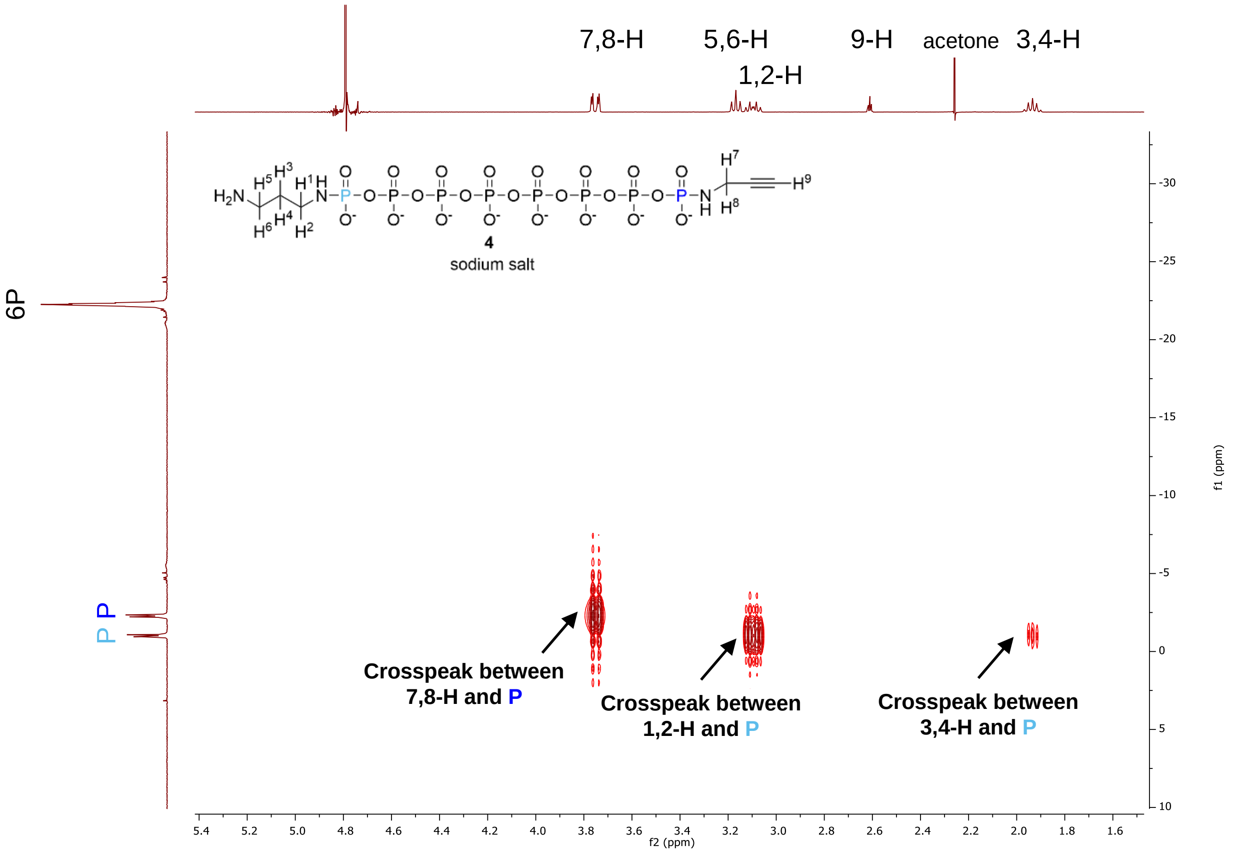

**Figure SI-1:** 2D $^1$H,$^{31}$P-HMBC spectrum of **4** to assign the phosphor signals. Assignment of proton signals was performed with $^1$H,$^1$H-COSY, $^1$H,$^{13}$C-HSQC and $^1$H,$^{13}$C-HMBC spectrum.

##### 1.5 Synthesis of sulfo-cyanin3-polyP$_8$-alkyne (5)

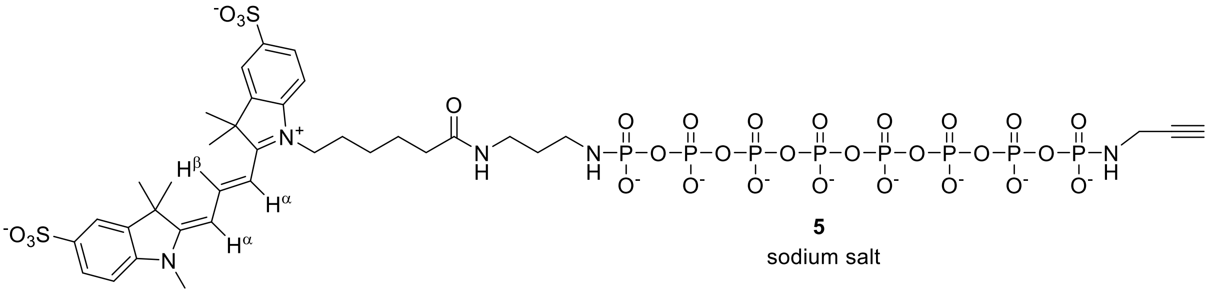

**5**
sodium salt

**4** (3.3 mg, 3.6 µmol, 1.0 eq.) was dissolved in 0.1 M $NaHCO_3$ buffer (450 µL) and was added to sulfo-cyanin3-NHS-ester (5.4 mg, 7.3 µmol, 2.0 eq.). The reaction mixture was stirred at r.t. for 3.5 h before it was diluted with water (20 mL) and directly injected to strong anion exchange chromatography system for purification (Q Sepharose® Fast Flow, increasing concentrations of 1 M $NaClO_4$ in $H_2O$). Fractions eluted with 12-28% aqueous $NaClO_4$ were combined and concentrated by lyophilization. Precipitation in cold $NaClO_4$ acetone solution (0.5 M, 35 mL), washing with cold acetone (3 × 15 mL) and drying afforded **5** (2.3 mg, 1.5 µmol, 42%) as a pink solid.

**$^1$H-NMR** (700 MHz, $D_2O$) δ = 8.62 (dd, *J* = 13.5, 13.5 Hz, 1H, **H**-β-Sulfo-Cy3), 7.95 (d, *J* = 1.7 Hz, 2H, 2x Ar-**H**), 7.91 (dd, *J* = 8.4, 1.8 Hz, 1H, Ar-**H**), 7.90 (dd, *J* = 8.4, 1.8 Hz, 1H, Ar-**H**), 7.45 (d, *J* = 7.9 Hz, 1H, Ar-**H**), 7.45 (d, *J* = 7.9 Hz, 1H, Ar-**H**), 6.46 (d, *J* = 13.9 Hz, 1H, **H**-α-Sulfo-Cy3), 6.44 (d, *J* = 13.9 Hz, 1H, **H**-α-Sulfo-Cy3), 4.18 (t, *J* = 7.5 Hz, 2H, Sulfo-Cy3-$N^+$-C**$H_2$**), 3.76 (dd, *J* = 10.6, 2.5 Hz, 2H, P-NH-C**$H_2$**-C≡C), 3.70 (s, 3H, Sulfo-Cy3-N-C**$H_3$**), 3.39 (s, N-**H**), 3.39 (s, N-**H**), 3.27 (t, *J* = 6.8 Hz, 2H, CO-NH-C**$H_2$**), 2.99 (dt, *J* = 9.7, 7.0 Hz, 2H, P-NH-C**$H_2$**), 2.62 (dt, *J* = 3.0, 1.5 Hz, 1H, C≡C-**H**), 2.31 (t, *J* = 7.4 Hz, 2H, NH-CO-C**$H_2$**), 1.90 (tt, *J* = 7.7, 7.7 Hz, 2H, P-NH-$CH_2$-C**$H_2$**), 1.83 (s, 6H, 2x C**$H_3$**), 1.82 (s, 6H, 2x C**$H_3$**), 1.75 – 1.67 (m, 4H, 2x Sulfo-Cy3-C**$H_2$**), 1.51 – 1.45 (m, 2H, Sulfo-Cy3-C**$H_2$**) ppm. **$^{31}$P{$^1$H}-NMR** (283 MHz, $D_2O$) δ = -0.39 – -0.72 (m, 1P), -2.04 – 2.42 (m, 1P), -21.55 – -22.45 (m, 6P) ppm. **$^{13}$C-NMR** (176 MHz, $D_2O$) δ = 176.7, 176.6, 176.0, 152.2, 145.2, 144.5, 141.8, 141.7, 139.2, 139.2, 126.6, 119.8, 119.7, 111.5, 111.3, 103.6, 103.5, 83.3, 71.5, 49.4, 49.3, 44.2, 39.1, 37.0, 35.7, 31.3, 31.1, 30.4 (d, *J* = 8.2 Hz), 27.2, 27.0, 26.6, 25.7, 25.2 ppm. **HRMS CE-ESI** calc. for $C_{36}H_{53}N_5O_{30}P_8S_2$ $[M-2H]^{2-}$: 673.5064, found: 673.5064.

#### 1.6 Synthesis of FRET-polyP$_8$ (6)

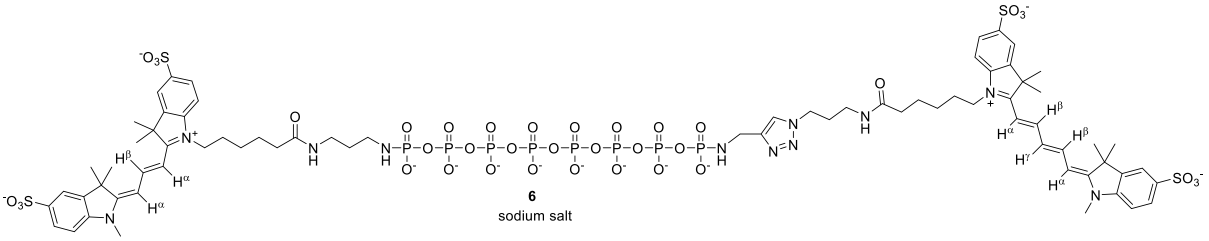

**5** (2.3 mg, 1.49 µmol, 1.0 eq.) was dissolved in 200 mM triethylammonium acetate buffer (pH 7, 1.6 mL). Sulfo-cyanin-5-azide (1.73 mg, 2.27 µmol, 1.5 eq.) was dissolved in water (220 µL) and was added to the mixture. The solution was degassed with an argon stream for 5 min. Subsequently, 20 mM $CuSO_4$ • 5 $H_2O$ solution (37 µL, 0.74 µmol, 0.5 eq.) and 50 mM THPTA solution (74 µL, 3.7 µmol, 2.5 eq.) were premixed and added. Then sodium ascorbate (3.0 mg, 15 µmol, 10 eq.) was added and the reaction mixture was stirred for 3.5 h at room temperature. The mixture was diluted with water (20 mL) and directly injected to strong anion exchange chromatography system for purification (Q Sepharose® Fast Flow, increasing concentrations of 1 M $NaClO_4$ in $H_2O$. Fractions eluted with 26-43% aqueous $NaClO_4$ were combined and concentrated by lyophilization. Precipitation in cold $NaClO_4$ acetone solution (0.5 M, 35 mL), washing with cold acetone (2 × 15 mL) and drying afforded a purple solid. The solid was dissolved in distilled $H_2O$ (1 mL) and chelex (2.5 mg) was added. After stirring gently for 30 min, the mixture was filtered through a syringe filter. Addition of chelex, stirring and filtration were repeated twice before the sample was lyophilized. **6** (1.2 mg, 0.52 µmol, 35%) was obtained as a purple solid.

**$^1$H-NMR** (700 MHz, $D_2O$) δ = 8.46 (dd, *J* = 13.4, 13.4 Hz, 1H, **H**-β-Sulfo-Cy3), 8.12 – 8.05 (m, 1H, Ar-**H**), 7.97 (dd, *J* = 12.7, 12.7 Hz, 1H, **H**-β-Sulfo-Cy5), 7.95 (dd, *J* = 12.8, 12.8 Hz, 1H, **H**-β-Sulfo-Cy5), 7.89 – 7.81 (m, 7H, 7x Ar-**H**), 7.41 (d, *J* = 8.7 Hz, 1H, Ar-**H**), 7.40 (d, *J* = 8.3 Hz, 1H, Ar-**H**), 7.38 (d, *J* = 8,1 Hz, 1H, Ar-**H**), 7.36 (d, *J* = 8.3 Hz, 1H, Ar-**H**), 6.55 (dd, *J* = 12.4, 12.4 Hz, 1H, **H**-γ-Sulfo-Cy5), 6.30 (d, *J* = 13.5 Hz, 1H, **H**-α-Sulfo-Cy3/5), 6.28 (d, *J* = 13.3 Hz, 1H, **H**-α-Sulfo-Cy3/5), 6.25 (d, *J* = 13.6 Hz, 1H, **H**-α-Sulfo-Cy3/5), 6.22 (d, *J* = 13.8 Hz, 1H, **H**-α-Sulfo-Cy3/5), 4.42 (t, *J* = 6.9 Hz, N(triazole)-C**H$_2$**), 4.25 (d, *J* = 10.2 Hz, 2H, P-NH-C**H$_2$**-triazole-SulfoCy5), 4.10 (t, *J* = 7.2 Hz, 2H, Sulfo-Cy3/5-N$^+$-C**H$_2$**), 4.00 (t, *J* = 7.8 Hz, 2H, Sulfo-Cy3/5-N$^+$-C**H$_2$**), 3.62 (s, 3H, Sulfo-Cy3/5-N-C**H$_3$**), 3.60 (s, 3H, Sulfo-Cy3/5-N-C**H$_3$**), 3.39 (s, 2H, 2x N-**H**), 3.39 (s, 2H, 2x N-**H**), 3.28 (t, *J* = 6.7 Hz, 2H, CO-NH-C**H$_2$**), 3.16 (t, *J* = 6.6 Hz, 2H, CO-

NH-C**H2**), 3.01 (dt, $J$ = 10.0, 7.1 Hz, 2H, P-NH-C**H2**-Linker-Sulfo-Cy3), 2.29 (t, $J$ = 7.4 Hz, 2H, NH-CO-C**H2**), 2.23 (t, $J$ = 7.2 Hz, 2H, NH-CO-C**H2**), 2.10 (tt, $J$ = 6.8, 6.8 Hz, 2H, N-CH2-C**H2**), 1.87 – 1.30 (m, 38H, 7x C**H2**, 8x C**H3**) ppm. **$^{31}$P{$^{1}$H}-NMR** (283 MHz, $D_2O$) δ = -0.44 – -0.71 (m, 1P), -1.66 – 1.97 (m, 1P), -21.51 – -22.48 (m, 6P) ppm. **HRMS CE-ESI** calc. for $C_{71}H_{96}N_{11}O_{37}P_8S_4$ $[M-3H]^{3-}$: 690.0923, found: 690.0917.

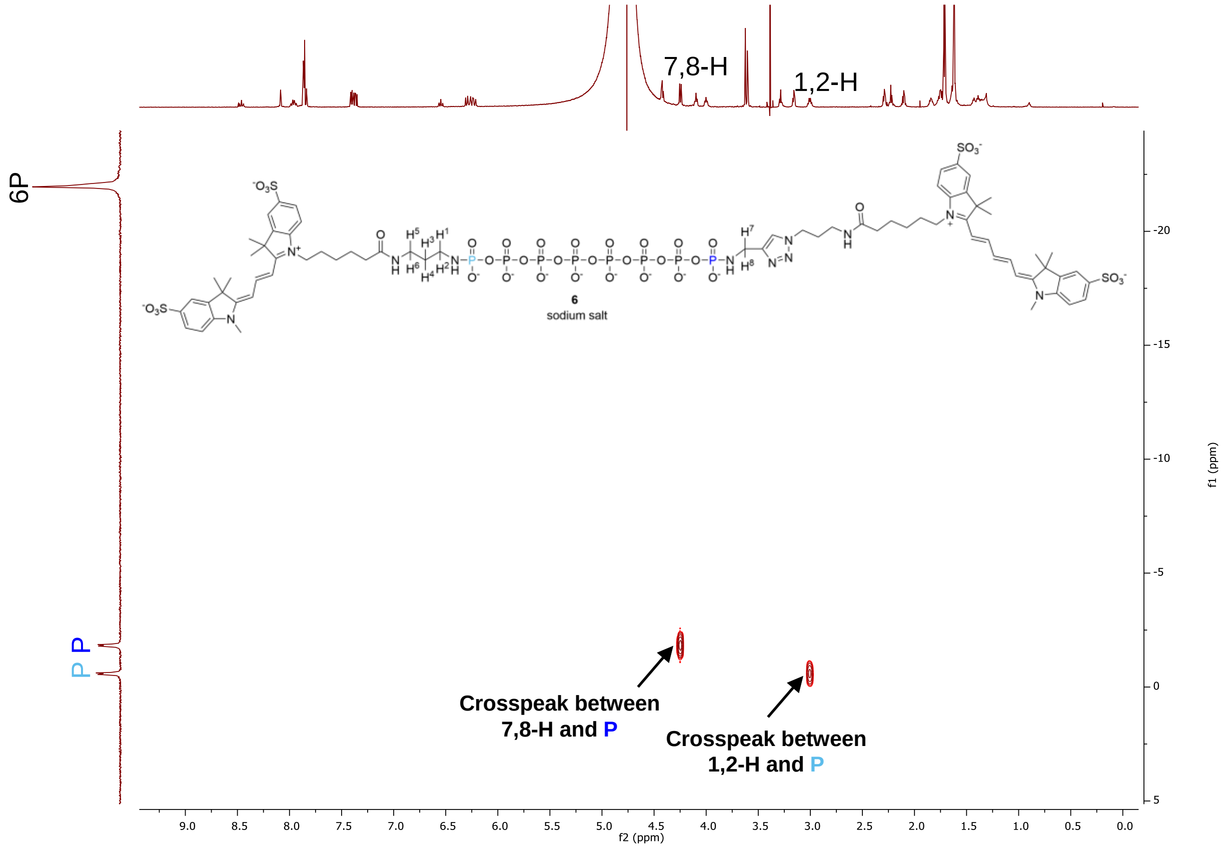

**Figure SI-2:** 2D $^{1}$H,$^{31}$P-HMBC spectrum of **6.**

#### 2. UPLC-MS Analysis

UPLC-MS analysis was performed on a UHPLC (Agilent 1290 infinity II) coupled to a qTOF (Agilent 6546). Chromatographic separation was achieved using gradient elution on a C18 column (AdvanceBio peptide plus 2.1mm x 150 mm) at 20 ˚C. Mobile phase consisted of 10 mM ammonium formate (pH 8.5) (eluent A) and 100% acetonitrile (eluent B). Chromatograms of **5** and **6** are presented in Figure SI-3 and SI-4, respectively.

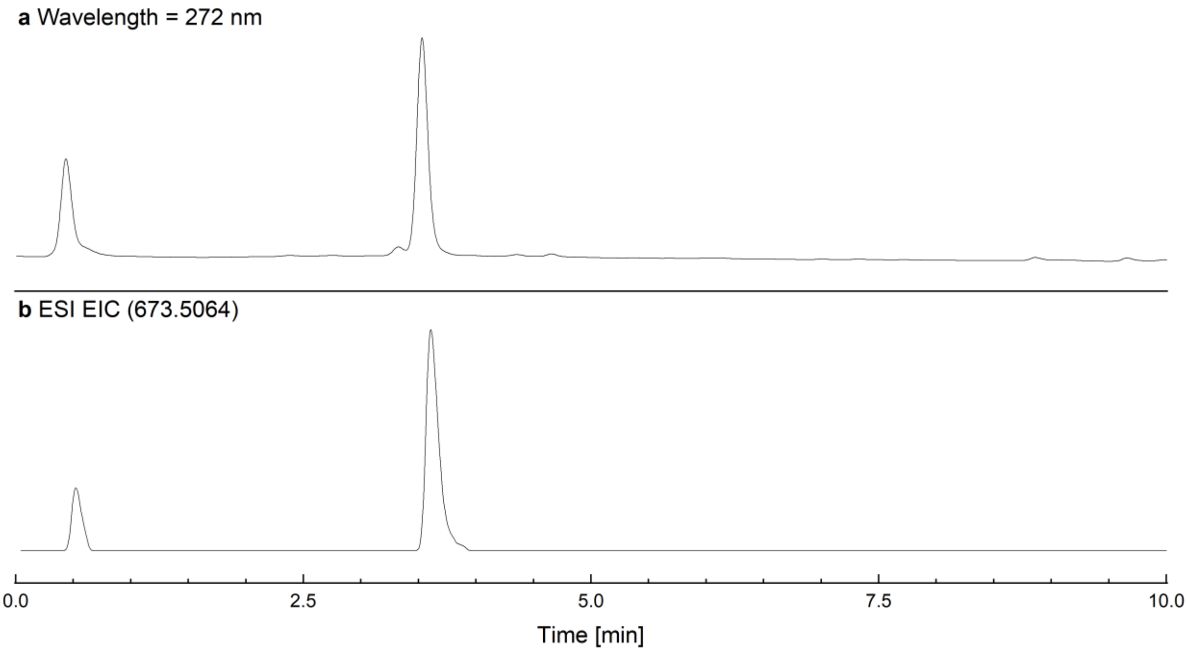

**Figure SI-3:** UPLC-MS-chromatogram of **5**. Run commenced with a linear gradient from 2% B to 40% B in 8 min, followed by a linear gradient to 80% B in 2 min with a flow rate of 0.5 mL/min. **a,** 272 nm UV detection. **b,** Extracted ion chromatogram (EIC) of **5** (m/z=673.5064 with z=2).

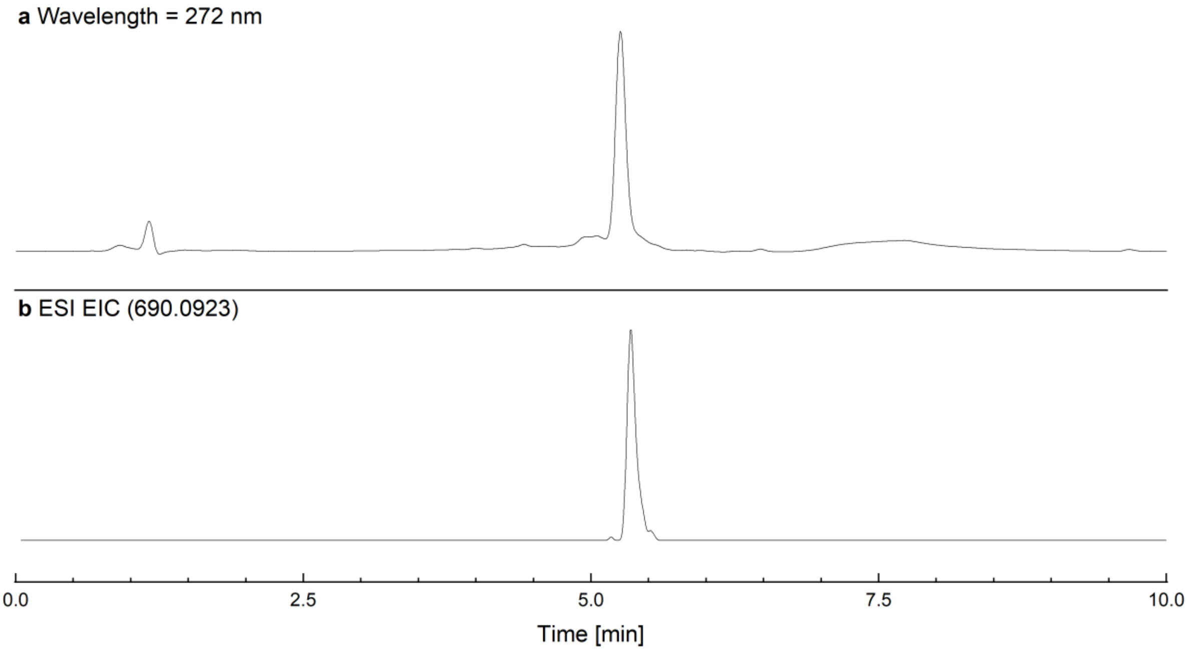

**Figure SI-4:** UPLC-MS-chromatogram of **6**. Run conducted with a linear gradient from 10% B to 40% B in 10 min with a flow rate of 0.4 mL/min. **a,** 272 nm UV detection. **b,** Extracted ion chromatogram of **6** (m/z=690.0923 with z=3).

#### 3. Isolation of Yeast Cytosolic Extract

The generation of the yeast strains *ppx1∆ppn1∆ppn2∆*, *ppx1∆ppn1∆ppn2∆vtc4∆*, and *ppx1∆ppn1∆ppn2∆ddp1∆*, were previously described.[4,5] Yeast cytosolic extract isolation protocol was adapted from Schmidt *et al*. (2010).[6] BY4742 yeast strains were grown in YPD medium (1% (w/v) yeast extract, 2% (w/v) bacto peptone, 2% (w/v) glucose) overnight at 30 °C and 130 rpm to an $OD_{600}$ of 1. 100 mL yeast cells were harvested by centrifugation and washed with distilled water and pelleted again by centrifugation. Cell pellets were treated with DTT and 0.15 mg/mL zymolyase 20T to generate spheroblasts. Spheroblasts were washed with 1.2 M sorbitol buffer and pelleted by centrifugation. Spheroblast pellets were lysed in 1 mL lysis buffer (20 mM Tris/HCl, pH 7.2, 30 mM NaCl, 10 mM $MgCl_2$, 10% glycerol, 1 mg/mL BSA, 2 mM PMSF). The steps following spheroblast lysis were performed on ice or in the cold room. 1 mL of sterile glass beads (0.25-0.5 mm diameter) were added onto samples and cells were opened by FastPrep-24 5G homogenizer at 4 m/sec for 20 s. The cell homogenate was centrifuged for 10 min at 2,500 × g and 4 °C and the supernatant was collected into a pre-cooled tube. To remove the non-cytosolic cellular fractions, the supernatant was further centrifuged for 15 min at 16,000 × g and 4 °C. The supernatant containing yeast cytosolic extracts was collected and the protein concentration was determined by Bradford Assay.[7] The cytosolic extracts were snap frozen in liquid nitrogen and stored at −80 °C.

#### 4. FRET Endopolyphosphatase Activity Assay

##### 4.1 With Isolated Proteins

A 3 µM working concentration was prepared from a 665 µM stock solution of FRET-polyP8 (**6**), whose concentration was determined by UV/Vis (Table SI-1). **6** (100 nM end concentration) in the corresponding 1× reaction buffer (Table SI-3) was supplemented with 2 µg of the corresponding enzyme. The total reaction volume was 60 µL. The samples were either incubated in a thermoshaker (30 min, 37 °C, 450 rpm) before being transferred to a 96-well plate for fluorescence spectrum measurement, or directly incubated in a plate reader (90 min, 37 °C, orbital shaking) to monitor fluorescence over time, with measurements taken every 5 min. Enzymes (GST-PPX from E. coli, His-DDP1 from *S. cerevisiae* and human GST-NUDT3, GST-

NUDT3E70A, GST-NUDT10, GST-NUDT11 were heterologously expressed and purified.[8,9]

**Table SI-3:** Composition of buffer solutions for FRET endopolyphosphatase activity assay.

| enzyme | concentration | buffer |
|---|---|---|
| **DDP1** / **PPX** | 1× | 25 mM HEPES, 50 mM NaCl, 10 mM $MgSO_4$, pH 6.8 |
| **NUDT3** / **NUDT3Δ** / **NUDT10** / **NUDT11** | 10× | without ion: 200 mM Tris pH 7.5, 1 M $NH_4Ac$ |
| | 10× | $Mg^{2+}$: 200 mM Tris pH 7.5, 1 M $NH_4Ac$, 50 mM $MgCl_2$ |
| | 10× | $Zn^{2+}$: 200 mM Tris pH 7.5, 1 M $NH_4Ac$, 50 mM $ZnSO_4$ x 7 $H_2O$ |

##### 4.2 With Yeast Extracts

The reaction mixture was prepared by sequentially adding lysis buffer (20 mM Tris pH 7.4, 30 mM NaCl, 10 mM $MgCl_2$, 10% glycerol, 2 mM PMSF), $MnSO_4$ (if applicable), **6** and yeast extract (1 mg) in a 96-well plate. The final concentrations in the 60 µL reaction volume were 100 nM **6** and 10 mM $MnSO_4$. Fluorescence specta were recorded at 5 min intervals over a 90 min period at 37 °C (Table SI-2).

#### 5. FRET Inhibition Assay of DDP1

A solution of FRET-polyP8 (**6**) at a final concentration of 100 nM in 1× reaction buffer (25 mM HEPES, 50 mM NaCl, 10 mM $MgSO_4$, pH 6.8) was first supplemented with the corresponding inhibitor solution followed by the addition of 2 µg of DDP1 or no enzyme (serving as decomposition control). The samples were incubated in a thermoshaker (30 min, 37 °C, 450 rpm) and then transferred to a 96-well plate for fluorescence spectrum analysis.

#### 5.1 Stability Against Different Potential DDP1 Inhibitors

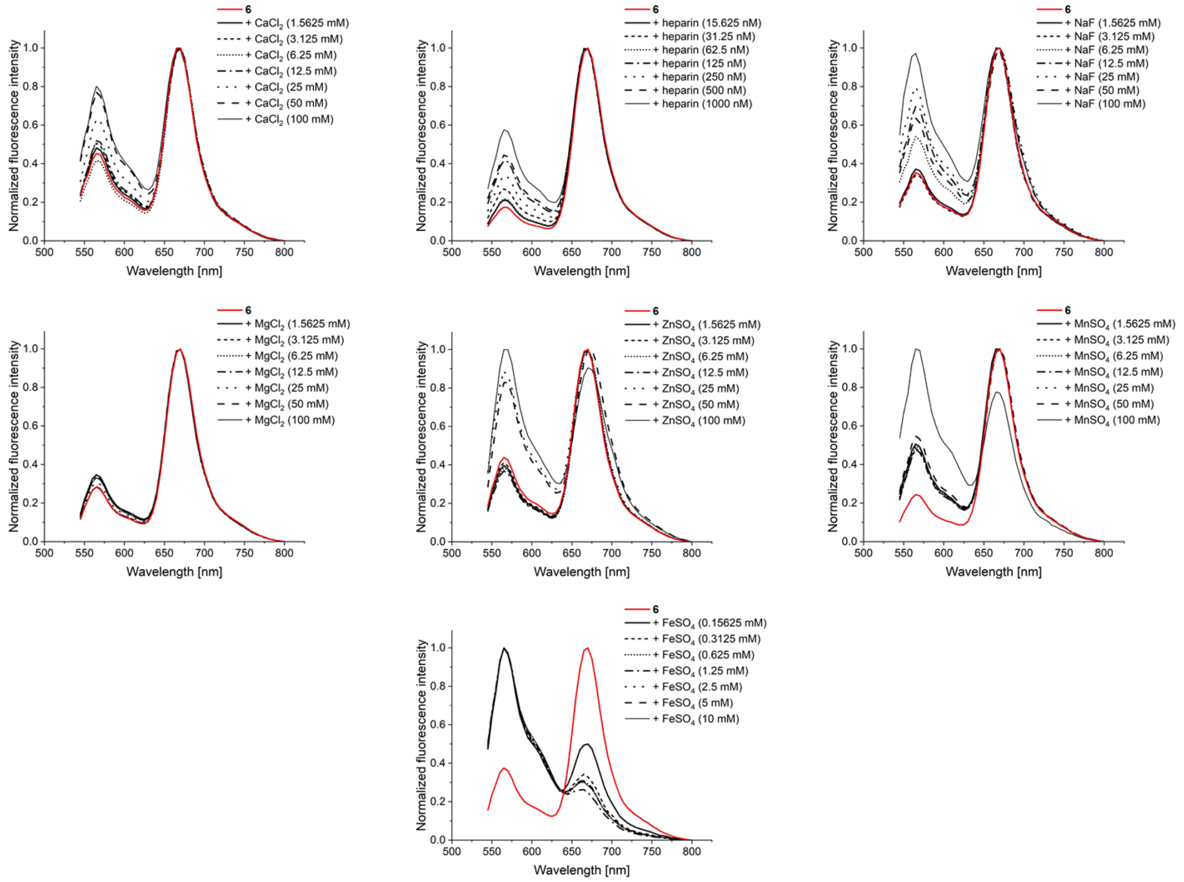

**Figure SI-5:** Normalized fluorescence intensity spectra of **6** (100 nM) in 1× reaction buffer (25 mM HEPES, 50 mM NaCl, 10 mM $MgSO_4$, pH 6.8) after incubation with different salt concentrations of different potential DDP1 inhibitors at 37 °C for 30 min (heparin sodium salt: 0-1,000 nM (calculated with average molecular weight of 17 kDa), $FeSO_4$: 0-10 mM, others: 0-100 mM). Excitation at $\lambda$ = 500 nm.

#### 5.2. DDP1 Inhibition Screening

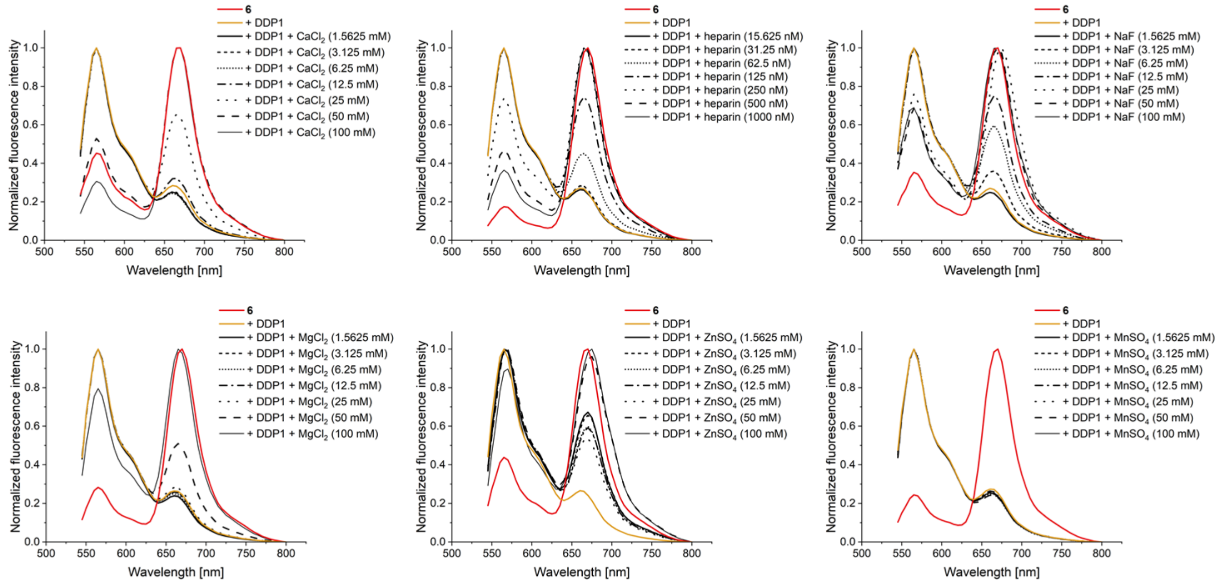

**Figure SI-6:** Normalized fluorescence intensity spectra of **6** (100 nM) in 1× reaction buffer (25 mM HEPES, 50 mM NaCl, 10 mM $MgSO_4$, pH 6.8) after incubation with DDP1 (2 µg) at 37 °C for 30 min in the presence of different potential inhibitors, tested across different concentrations. Excitation at λ = 500 nm.

### 6. PAGE Endopolyphosphatase Activity Assay in Yeast Extracts

The generation of the yeast strain *ppx1Δppn1Δppn2Δddp1Δ* was previously described.[4] To prepare protein extracts logarithmic growing yeast culture (20-50U of $OD_{600}$) were harvested by centrifugation at 5,000 × g for 3 min and washed once with ice-cold Milli-Q water. Proteins were extracted in ice-cold lysis buffer (50 mM Tris-HCl, pH 7.4, 100 mM NaCl with fresh added 5 mM DTT, and protease inhibitor mixture (SigmaAldrich, P8215)) by vortexing in the presence of acid washed glass beads for 5 min at 4 °C. The homogenates were centrifuged 15,000 × g for 5 min at 4 °C and the supernatants (protein extracts) were transferred to another tube and used immediately. Protein extracts (100 µg) were incubated in reaction buffer (25 mM Tris-HCl, pH 7.4, 100 mM NaCl, 1 mM DTT) alone or supplemented with 10 mM MgCl or 10 mM MnCl for 60 min at 37 °C. After the incubation, the reactions were resolved on a 30% polyacrylamide gel and polyP visualized by toluidine blue staining as previously described.[10]

### 8. NMR Spectra

1H-NMR (400 MHz, D2O)

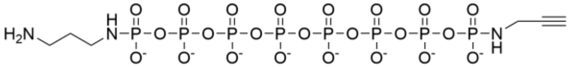

**4**
sodium salt

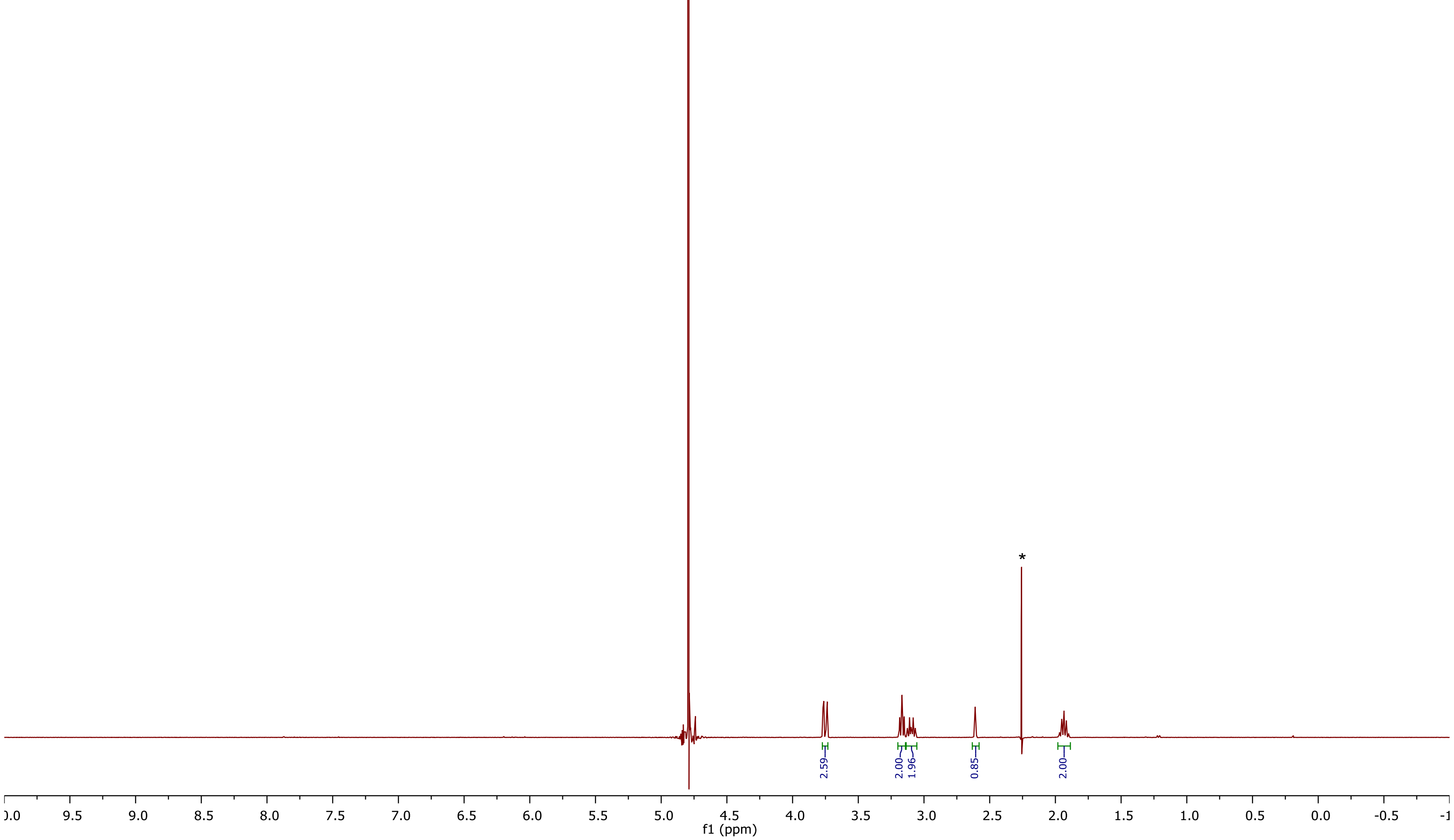

Residual amounts of acetone are marked with asterisks (*).

31P{1H}-NMR (162 MHz, D2O)

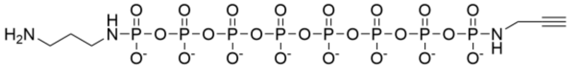

**4**

sodium salt

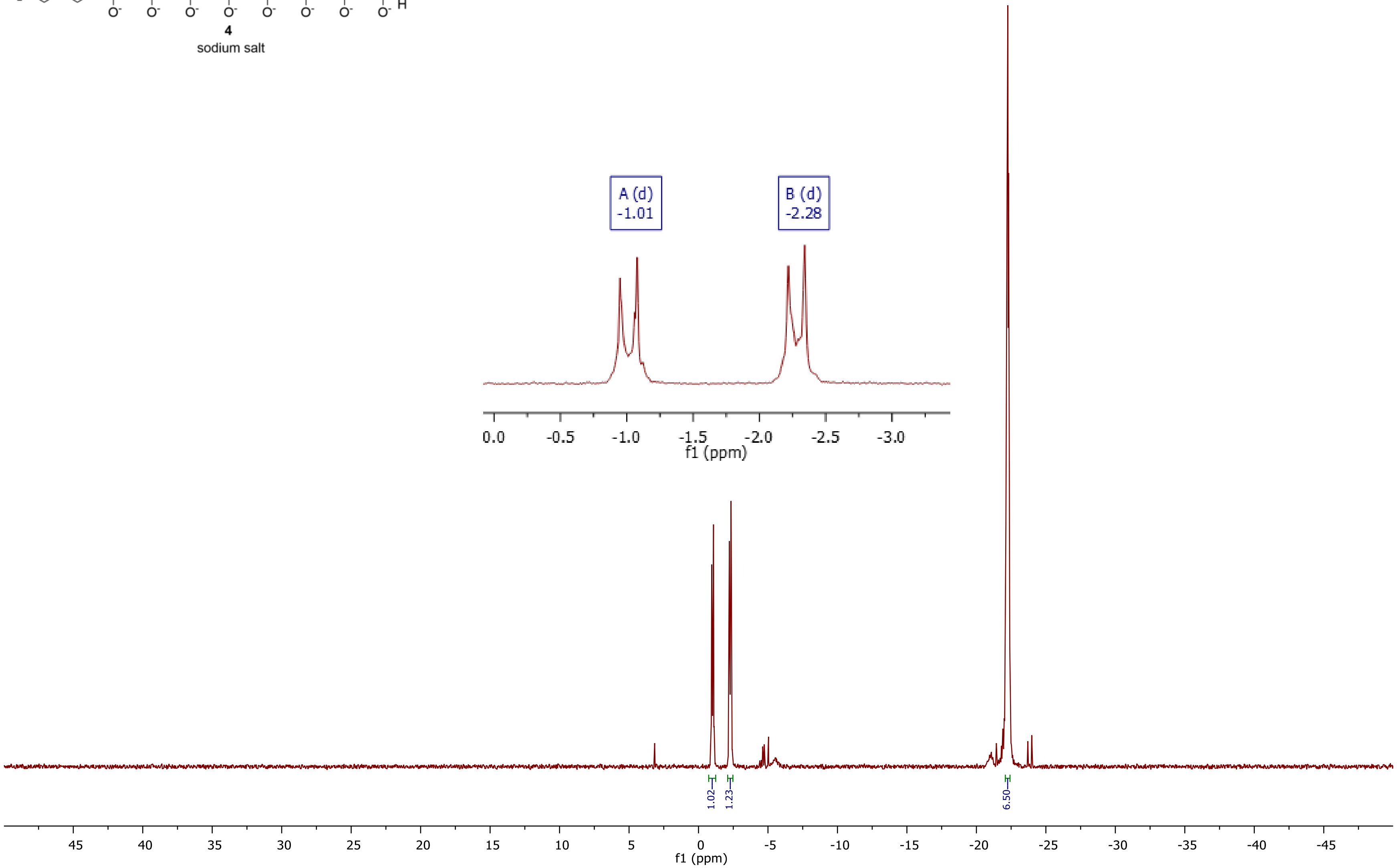

31P-NMR (162 MHz, D2O)

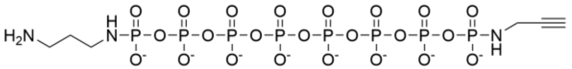

4

sodium salt

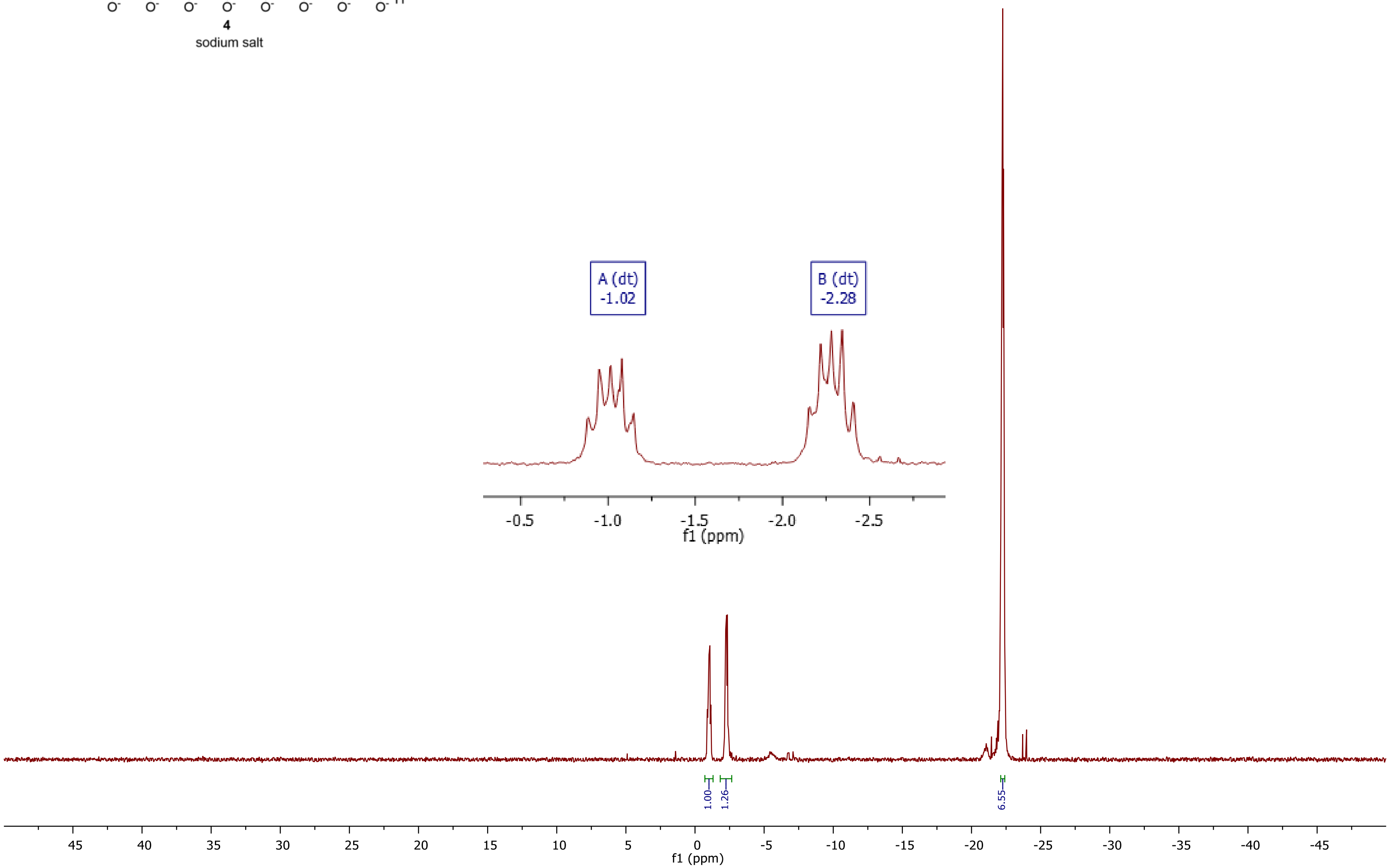

13C-NMR (101 MHz, D2O)

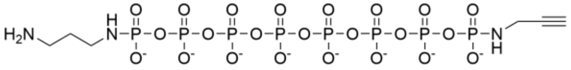

**4**

sodium salt

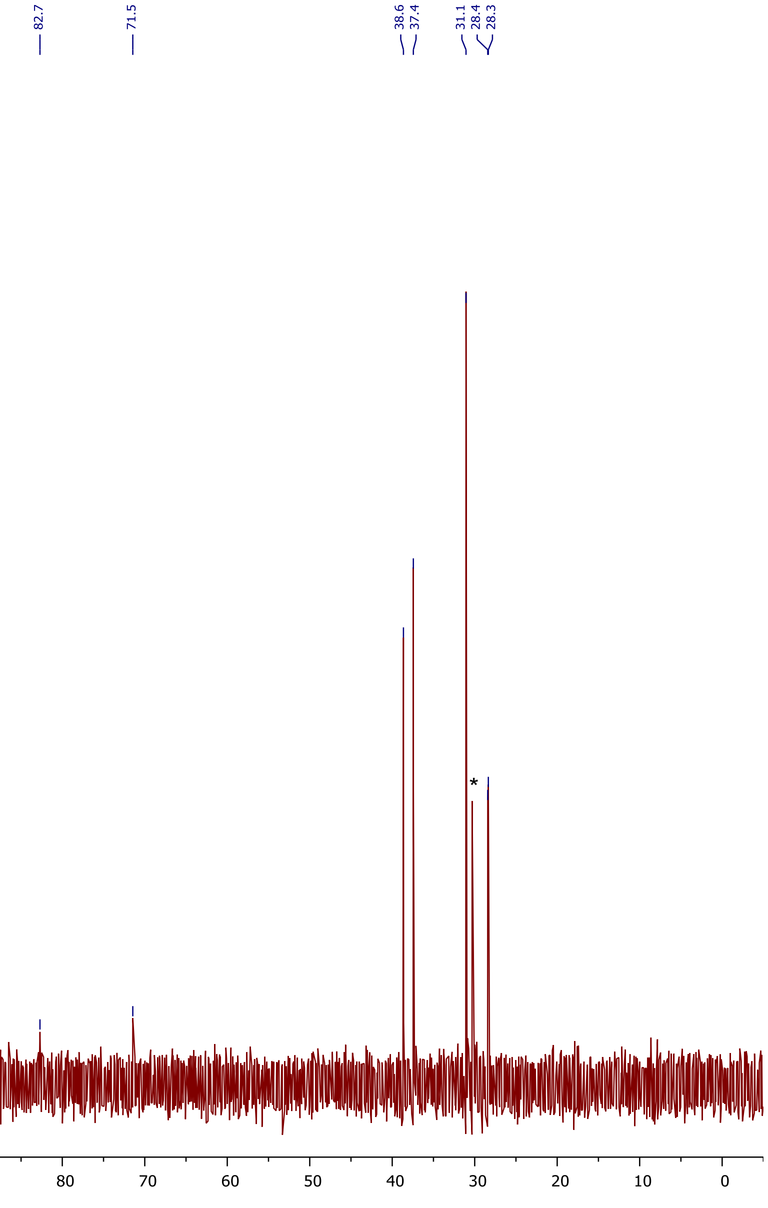

Residual amounts of acetone are marked with asterisks (*).

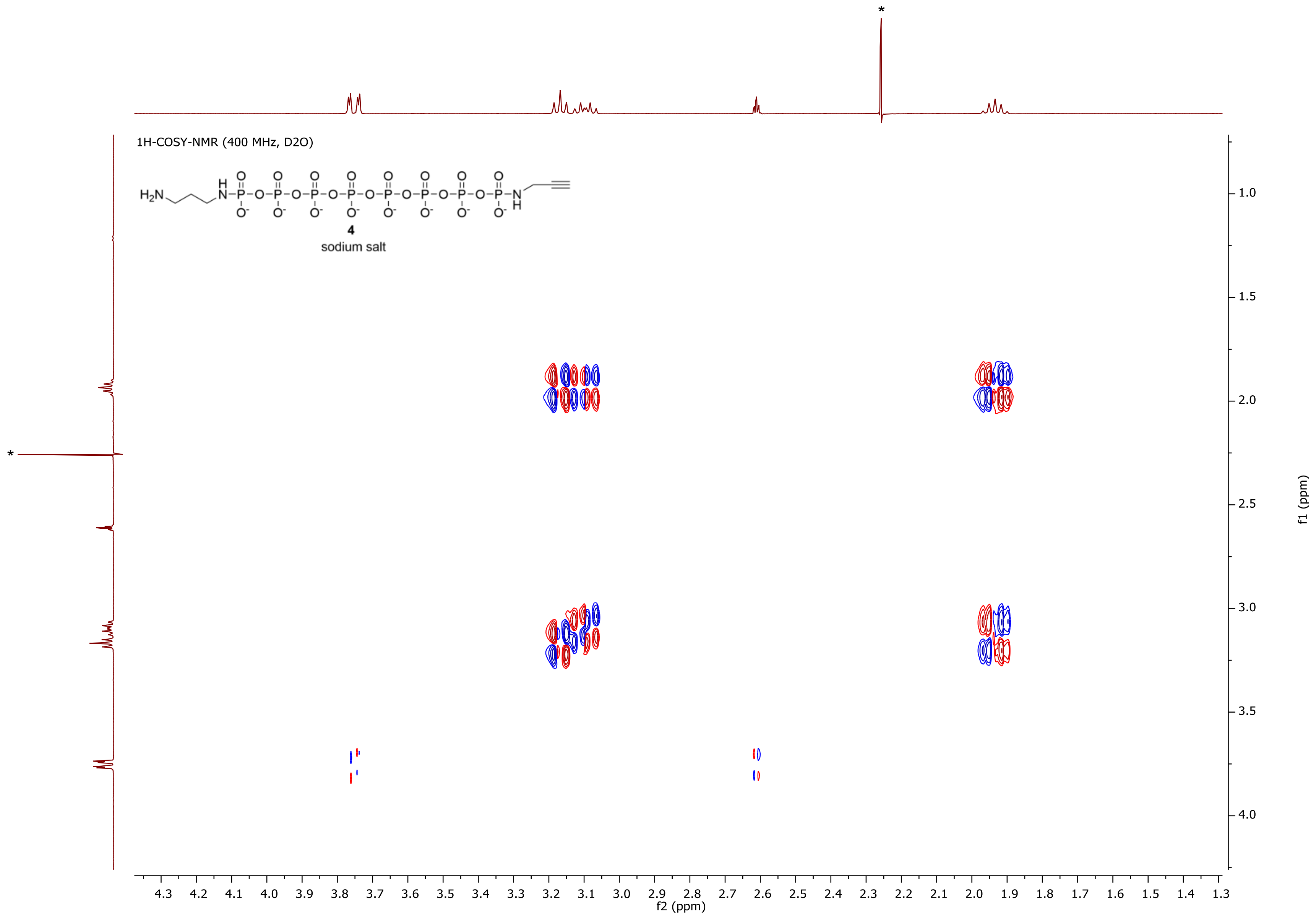

Residual amounts of acetone are marked with asterisks (*).

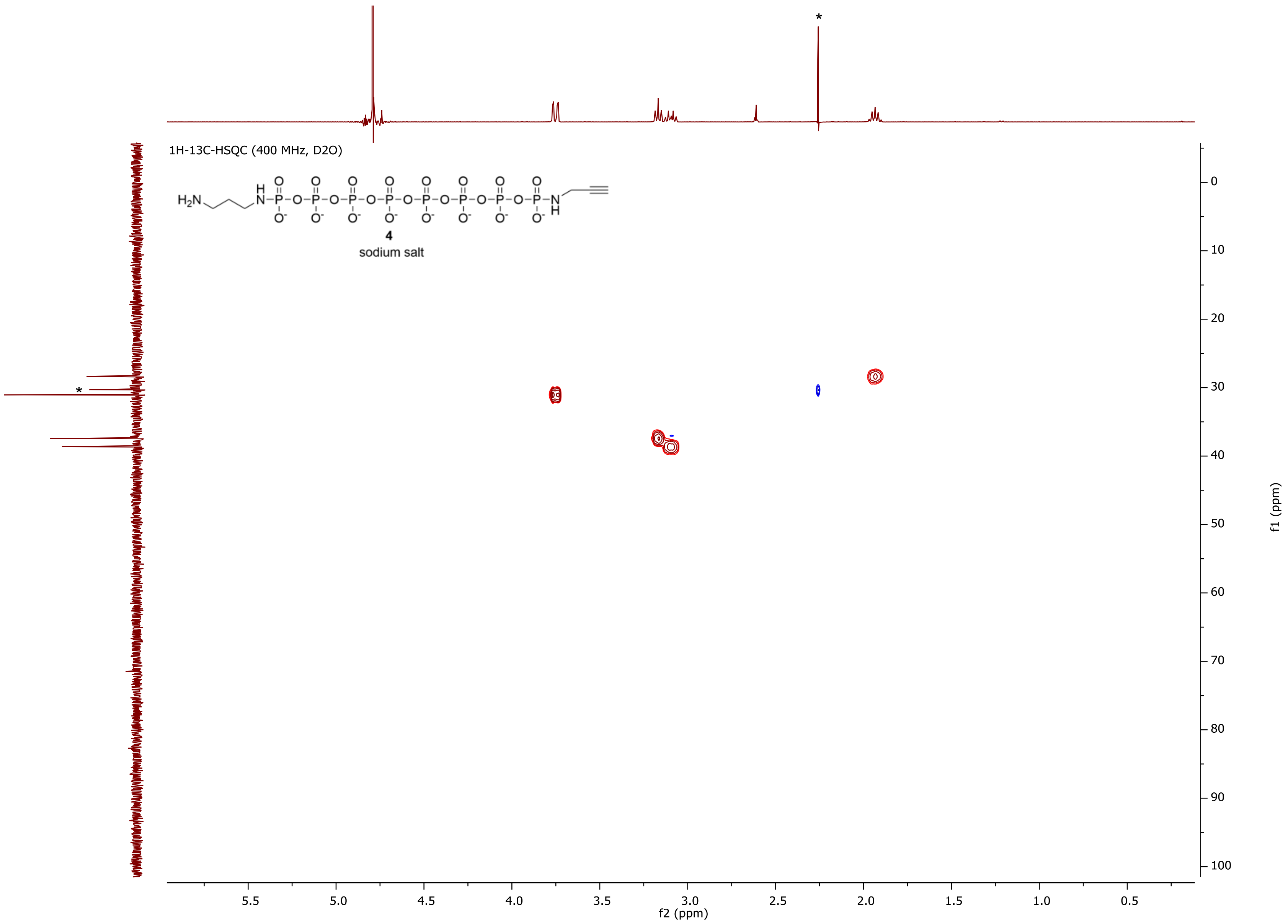

Residual amounts of acetone are marked with asterisks (*).

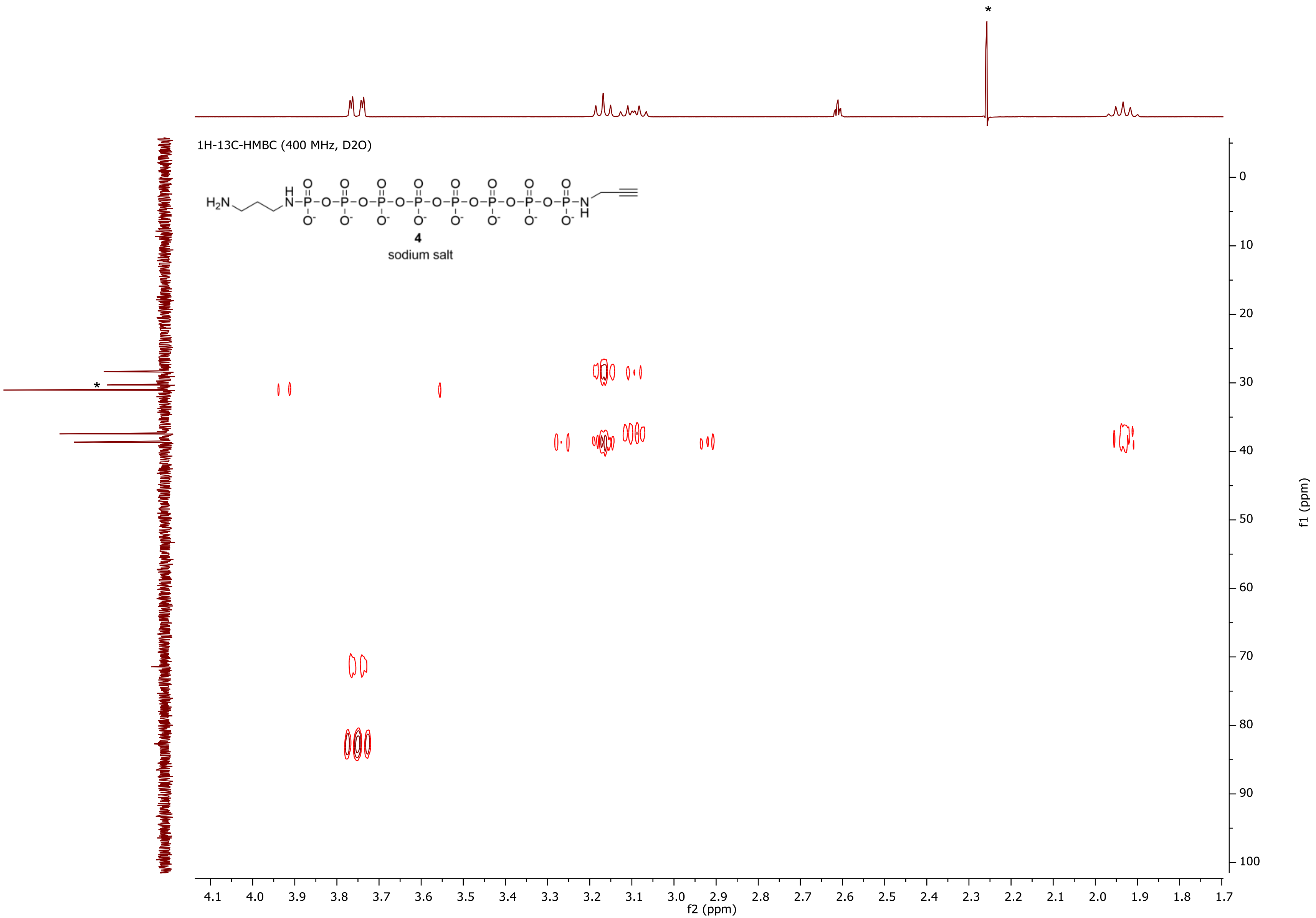

Residual amounts of acetone are marked with asterisks (*).

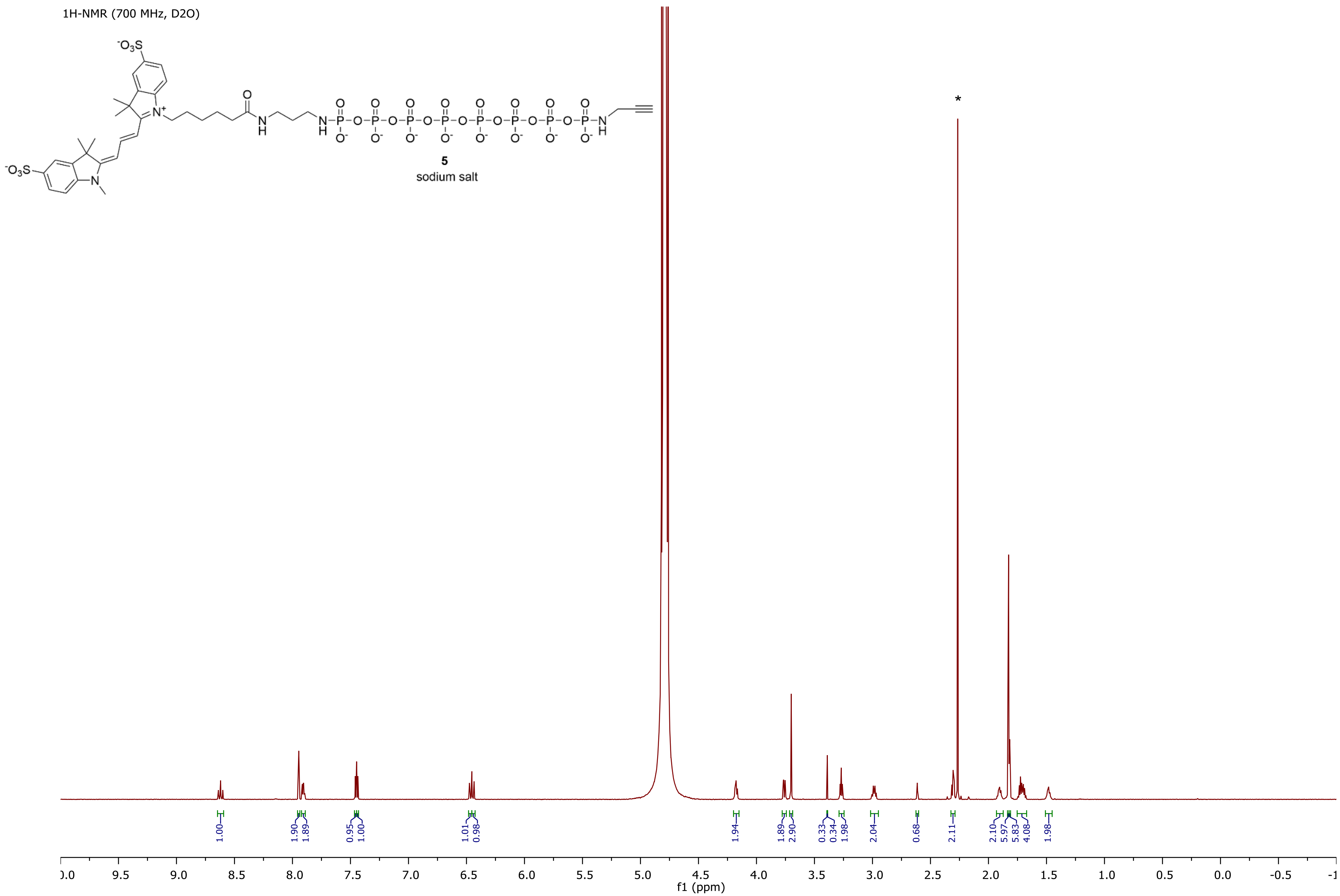

Residual amounts of acetone are marked with asterisks (*).

31P{1H}-NMR (283 MHz, D2O)

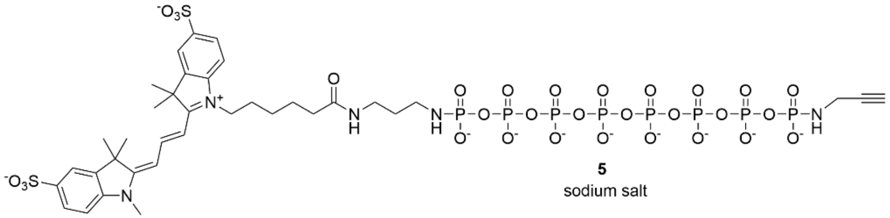

**5**
sodium salt

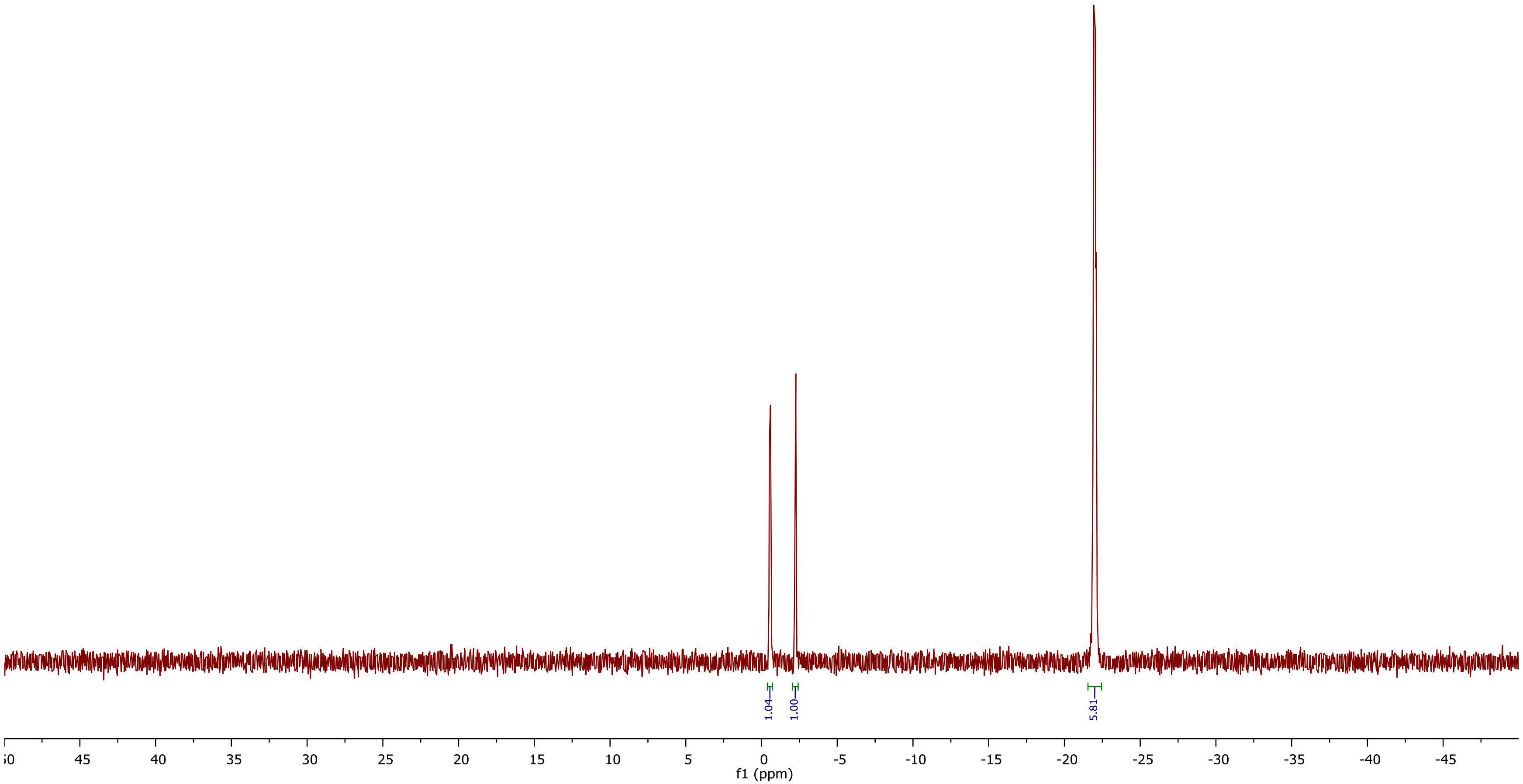

13C-NMR (176 MHz, D2O)

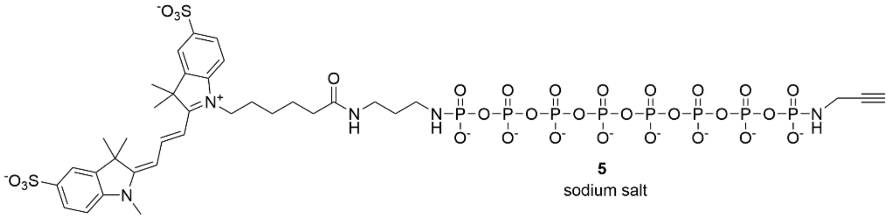

5
sodium salt

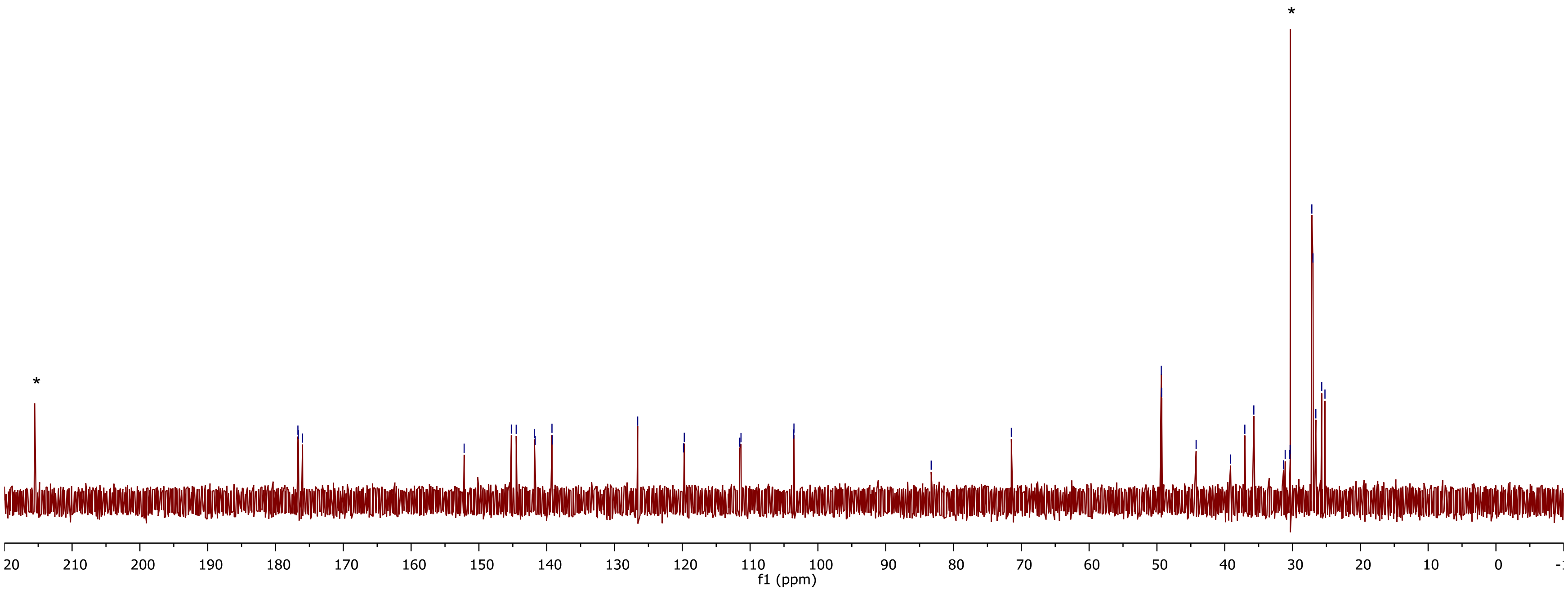

Residual amounts of acetone are marked with asterisks (*).

1H-NMR (700 MHz, D2O)

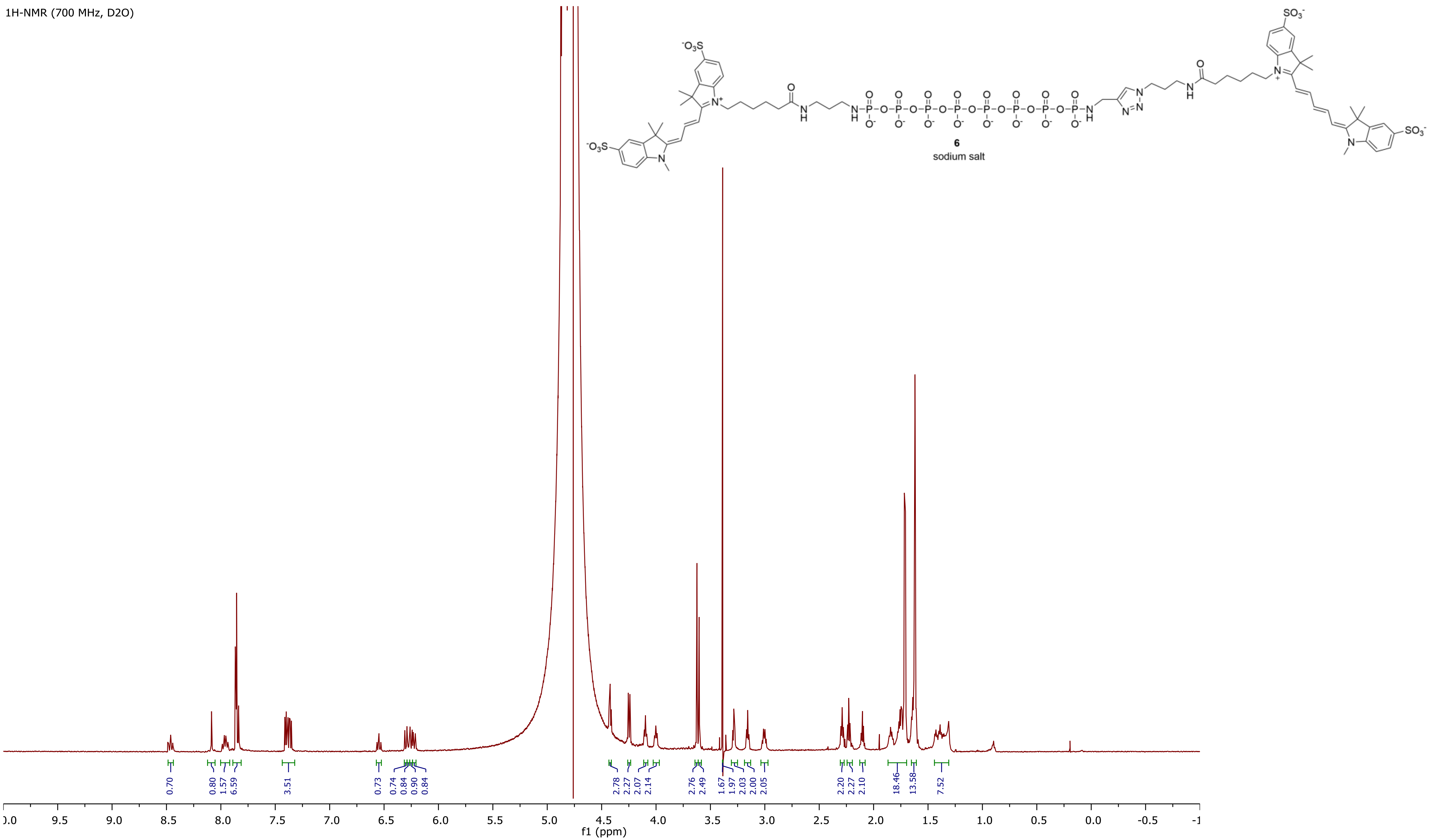

31P{1H}-NMR (283 MHz, D2O)

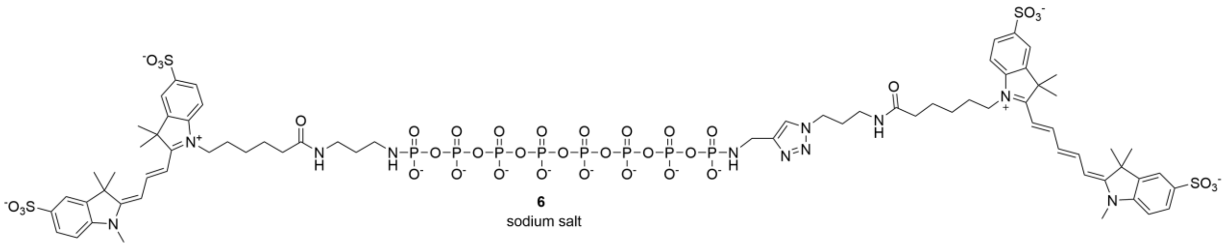

6
sodium salt

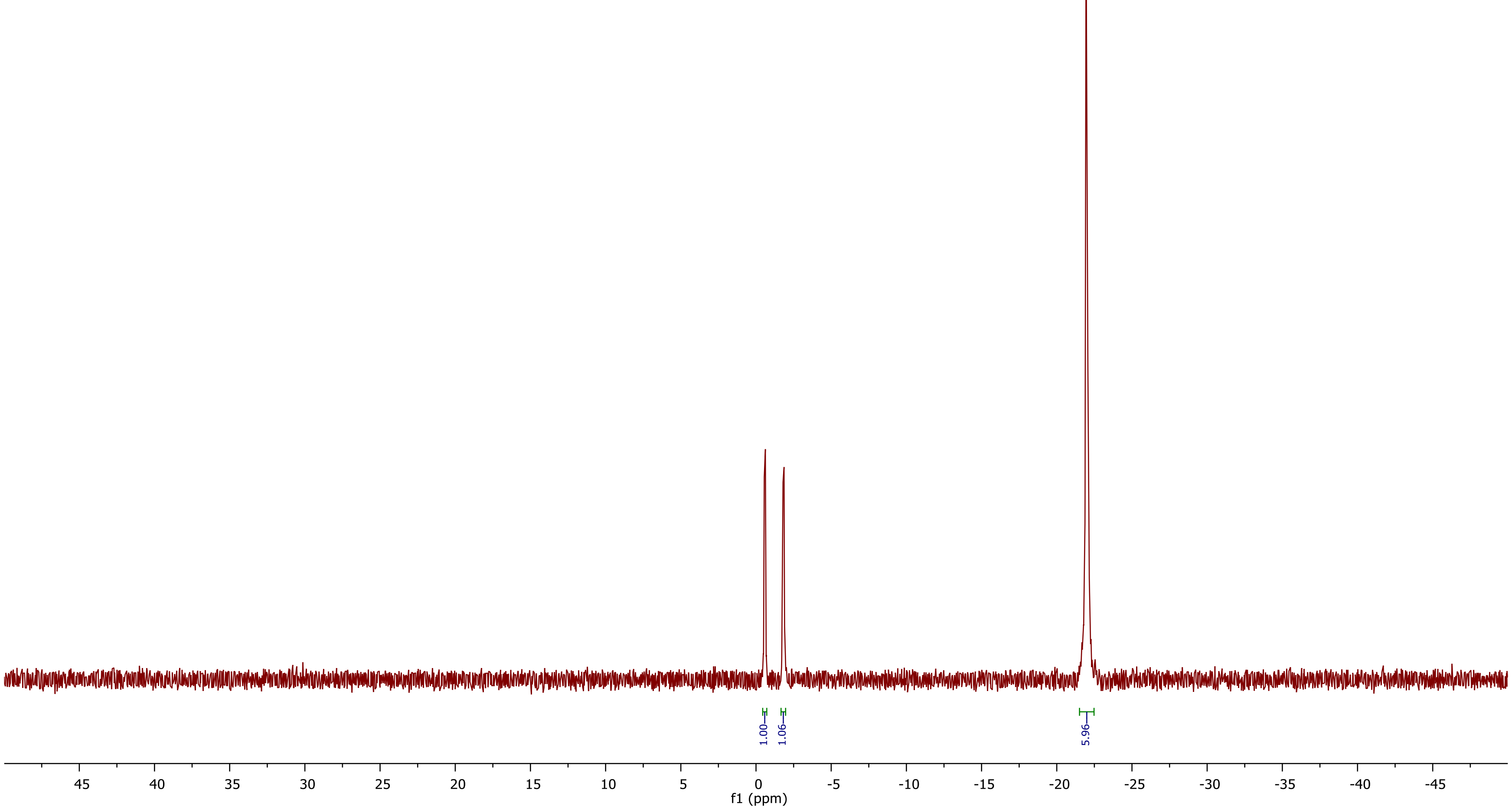
